# The Arabidopsis SHORTROOT network coordinates shoot apical meristem development with auxin dependent lateral organ initiation

**DOI:** 10.1101/2022.09.27.509828

**Authors:** Elmehdi Bahafid, Imke Bradtmöller, Ann Marlene Thies, Thi Thuy Oanh Nicole Nguyen, Crisanto Gutierrez, Bénédicte Desvoyes, Yvonne Stahl, Ikram Blilou, Rüdiger Simon

**Affiliations:** Institute for Developmental Genetics, Heinrich Heine University, Building 26.14.00.071, 40225 Düsseldorf, Germany; Centro de Biologia Molecular Severo Ochoa, CSIC-UAM, Cantoblanco, Madrid, Spain; Laboratory of Plant Cell and Developmental Biology, Division of Biological and Environmental Sciences and Engineering, 4700 King Abdullah University of Science and Technology, Ibn Al Haytham Bldg.2, room 3274, Thuwal 23955-6900, Kingdom of Saudi Arabia

## Abstract

Unlike animals, plants have the capacity to produce new organs post-embryonically throughout their entire life cycle. This is due to stem cells present in the shoot and the root apical meristems (SAM and RAM, respectively). In the SAM, stem cells are located in the central zone (CZ) where they divide slowly. Stem cell daughters are displaced laterally and enter the peripheral zone (PZ). Here, their mitotic activity increases, and lateral organ primordia (LOP) are formed. How the spatial arrangement of these different domains is initiated and controlled during SAM growth and development, and how sites of LOP are determined in the PZ is not yet completely understood.

In the RAM, the GRAS family transcription factor SHORTROOT (SHR) acts as a master regulator of signalling pathways that maintain the root stem cell niche and control formation of ground tissue layers. We hypothesized that SHR could perform a similar role in the SAM, and found that SHR, together with its target transcription factors SCARECROW (SCR), SCARECROW-LIKE23 (SCL23) and JACKDAW (JKD), controls shoot meristem size by regulating cell division rates, and promotes formation of lateral organs. SHR, SCR, SCL23 and JKD are expressed in very distinct patterns in the SAM. Where these expression domains overlap, they can physically interact to activate expression of the key cell cycle regulator *CYCLIND6;1* (*CYCD6;1*) and thereby promote the formation of new cell layers.

In the PZ, upregulation of *SHR* expression at sites of organ initiation depends on the phytohormone auxin, acting through the auxin response factor MONOPTEROS (MP) and auxin response elements in the *SHR* promoter. In the CZ, the SHR-target SCL23 physically interacts with WUS, a key regulator of stem cell maintenance, and both SCL23 and WUS expression are subject to negative feedback regulation from stem cells through the CLAVATA signalling pathway. Together, our findings illustrate how SHR-dependent transcription factor complexes act in different domains of the shoot meristem to mediate cell division and auxin dependent organ initiation in the PZ, and coordinate this activity with stem cell maintenance in the CZ of the SAM.

## Introduction

In multicellular organisms, stem cells are the source of all tissues and organs. In contrast to animals where organs are formed during embryogenesis, plants form organs only postembryonically, and they can continuously produce new organs throughout their life span. This process depends on the activity of pluripotent stem cells embedded in their shoot and root apical meristems (SAM and RAM, respectively).

In the SAM, stem cells are located in the central zone (CZ) at the tip of the meristem. Directly underneath lies the organizing center (OC), which is required for stem cell maintenance. The stem cells in the CZ have a low mitotic activity, but after cell division, descendants are continuously displaced towards the peripheral zone (PZ). Here, they start to divide faster and provide founder cells for lateral organ primordia (LOP) that will develop at the meristem flank (Reddy *et al*., 2004). How the communication between these distinct functional domains is coordinated to balance stem cell proliferation in the center with organ formation at the periphery of the SAM is still not completely understood. This is in contrast to the RAM where the gene networks regulating the spatial arrangement of the different tissue types are well known. In the RAM, a member of the plant-specific GRAS transcription factor (TF) family, SHORT-ROOT (SHR) has a key regulatory role in the maintenance of the root stem cell niche identity and tissue patterning (Pysh *et al*., 1999, Lee *et al*., 2008, Long *et al*., 2015b, Helariutta *et al*., 2000, Nakajima *et al*., 2001). Plants lacking functional SHR show a ‘short root’ phenotype, with roots carrying fewer ground tissue layers (Di Laurenzio *et al*., 1996, Scheres *et al*., 1995, Pysh et al., 1999). *SHR* is transcribed in the stele, and SHR protein moves one cell layer outward where it interacts with the GRAS domain TFs SCARECROW (SCR) and SCARECROW LIKE 23 (SCL23), and with the BIRD/INDETERMINATE DOMAIN TF JACKDAW (JKD) (Long et al., 2015b, Long *et al*., 2017, Nakajima et al., 2001, Cui *et al*., 2007). The resulting multimeric complex promotes asymmetric cell division (ACD) of the Cortex-Endodermis Initial/daughter (CEI/CEID) by directly activating the expression of the key cell cycle regulator *CYCLIND6;1* (*CYCD6;1*) to instruct development of distinct endodermis and cortex cell layers (Long *et al*., 2015a, Long et al., 2015b, Long et al., 2017, Di Laurenzio et al., 1996, Helariutta et al., 2000, Nakajima et al., 2001). SCR and SHR are required for the maintenance and specification of the QC, as *shr* and *scr* mutants show abnormal QC cells (Sabatini et al., 2002, Petersson et al., 2009)*. SHR*, *SCR* and *SCL23* are also expressed in leaves and the SAM, but their function was mainly studied in leaves (Wysocka-Diller *et al*., 2000, Gardiner *et al*., 2011, Wang *et al*., 2011, Cui *et al*., 2014). Here, we asked whether patterning in the SAM involves the SHR signaling pathway in a mechanism similar to the RAM..

In the SAM, stem cell maintenance depends on the homeodomain transcription factor WUSCHEL (WUS) (Laux *et al*., 1996, Zhang *et al*., 2017). *WUS* transcripts are limited to the OC, but WUS protein moves via plasmodesmata from the OC to the CZ to directly activate the transcription of the secreted signalling peptide CLAVATA3 (CLV3) in stem cells (Mayer *et al*., 1998, Yadav *et al*., 2011, Daum *et al*., 2014). In turn, CLV3 acts to limit *WUS* expression to the OC via a system of receptor-like kinases and co-receptors: CLAVATA1 (CLV1), CLAVATA2 (CLV2), CORYNE (CRN) and BARELY ANY MERISTEM1-3 (BAM1-3) (Bleckmann & Simon, 2009, Clark *et al*., 1997, Fletcher *et al*., 1999, Nimchuk *et al*., 2015, Ohyama *et al*., 2009). The resulting WUS-CLV3 negative feedback circuit controls stem cell homeostasis in the SAM (Brand *et al*., 2000, Schoof *et al*., 2000). Recently it was shown that WUS interacts in the OC with the GRAS family transcription factors HAIRY MERISTEM 1 and 2 (HAM1/2) to restrict CLV3 activation to the apical domain of the CZ (Han *et al*., 2020).

Formation of lateral organ primordia at the meristem flanks is preceded by establishment of a local maximum of the plant hormone auxin (Benková *et al*., 2003, Heisler *et al*., 2005, Vernoux *et al*., 2011a). This auxin maximum is generated by the auxin efflux carrier PIN-FORMED1 (PIN1) through polar auxin transport (Friml *et al*., 2004, Gälweiler *et al*., 1998). The transcriptional auxin read-out is carried out by the Auxin/Indole-3-acetic acid (Aux/IAA) repressor proteins that under low auxin conditions dimerize with auxin response factors (ARFs) and repress their activity (Mockaitis & Estelle, 2008). ARFs bind DNA at the promoters of their target genes via an auxin response element (AuxRE) (Boer *et al*., 2014). An increase in auxin levels triggers the formation of a complex between Aux/IAA and TIR1/AFB leading to the degradation of Aux/IAA and the release of the ARFs, which can now regulate expression of auxin response genes (Paque & Weijers, 2016). Auxin-dependent lateral organ initiation is mediated by AUXIN RESPONSE FACTOR 5/MONOPTEROS (ARF5/MP) (Berleth & Jürgens, 1993, Hardtke & Berleth, 1998). MP plays a crucial role in the initiation of flower primordia as the strong *mp* allele lacks roots and a weak allele cannot produce flowers and forms a naked inflorescence stem (Yamaguchi *et al*., 2013). MP directly induces the expression of its target gene *LEAFY (LFY),* a plant-specific transcription factor that plays a key role in flower primordia initiation and in primordium fate specification (Schultz & Haughn, 1991, Weigel *et al*., 1992, Yamaguchi et al., 2013, Wu *et al*., 2015). Transcript levels of *SHR* and *SCR* are strongly reduced in *mp* mutant embryos (Möller *et al*., 2017) suggesting that MP regulates expression of *SHR* and *SCR* during embryonic development.

We here started to unravel how organ initiation at the periphery is coordinated with stem cell behaviour in the center of the SAM. We show that SHR, SCR, SCL23 and JKD act in the SAM in different expression domains with complementary and overlapping patterns, and together, they cover all the functional domains of the SAM. Interestingly, the overlapping region of the four TFs coincides with the expression of *CYCLIND6;1 (CYCD6;1).* We find that mutants for genes in the SHR gene regulatory network show reduced meristem sizes and delay cell cycle progression due to an extended G1 phase. We demonstrate that auxin via MP promotes expression of *SHR* and *SCR* in the PZ, where SHR acts upstream of *LFY* to promote flower primordia formation. In the meristem centre, members of the SHR network directly interact with the CLV3-WUS circuit. Together, our study shows how mobile TFs of the SHR network establish a balance between stem cell maintenance and lateral organ initiation, by controlling cell division rates and cell fate in different domains within the SAM in an auxin dependent manner.

## Results

### *SHR* and *SCR* coordinate shoot meristem size and primordia initiation

In order to identify SHR and SCR regulatory roles in shoot meristem development, we first analyzed the phenotypes of null mutant alleles of *SHR* (*shr-2*) and *SCR* (*scr-3* and *scr-4*). All mutants displayed a small rosette, dwarfed shoot phenotype and initiated fewer flowers compared to wild type (WT) (Fig. 1A-D; Suppl Fig. 1A-D). To assess the function of *SHR* and *SCR* in the SAM, we measured meristem sizes and the average time interval between the initiation of successive lateral organs (plastochron). Plastochrons were increased in the mutants (*shr-2, scr-3* and *scr-4*) compared to WT, indicating a significant delay in the initiation of lateral organs in the mutants (Fig. 1M; Suppl Fig. 1A’-D’). Next, we determined meristem size using confocal microscopy by imaging the meristem of mutants and WT (Fig. 1E-H) and measuring the surface area of the SAM excluding organ primordia as a proxy for meristem size. We found that the meristem is significantly reduced in the mutants (Fig. 1N), and hypothesized that SHR and SCR may serve to coordinate meristem size with lateral organ primordia initiation during SAM development. To better understand the cause for the reduction in SAM size, we analyzed both cell number and area on the meristem surfaces. Analysis of average cell surface areas showed no differences between mutants and WT (Suppl Fig. 1E), however, cell numbers were reduced in all mutants (Fig. 1O). We next investigated if there were any significant differences in cell proliferation rate between WT, *shr-2*, *scr-3* and *scr-4* at the SAMs. We used the plant cell cycle marker *PlaCCI* (Desvoyes *et al*., 2020) (Fig. 1I-L), an imaging tool combining the three reporters *pCDT1a:CDT1a-eCFP*, *pHTR13:HTR13-mCherry* and *pCYCB1;1:NCYCB1;1-YFP*, which individually mark the G1, S + G2 and mitotic (M) phases, respectively. In shoot meristems (excluding organ primordia) of *shr-2*, *scr-3* and *scr-4*, the percentage of cells in G1 marked with CDT1a-CFP was significantly increased compared to WT (Fig. 1P; Suppl Fig. 1F), indicating delayed progression through the cell cycle. We conclude that *SHR* and *SCR* promote cell cycle progression in the meristem by controlling the G1 phase, which could account for the observed reduction in SAM size and delay in organ initiation in the PZ of *shr* and *scr* mutants.

**Fig. 1:**
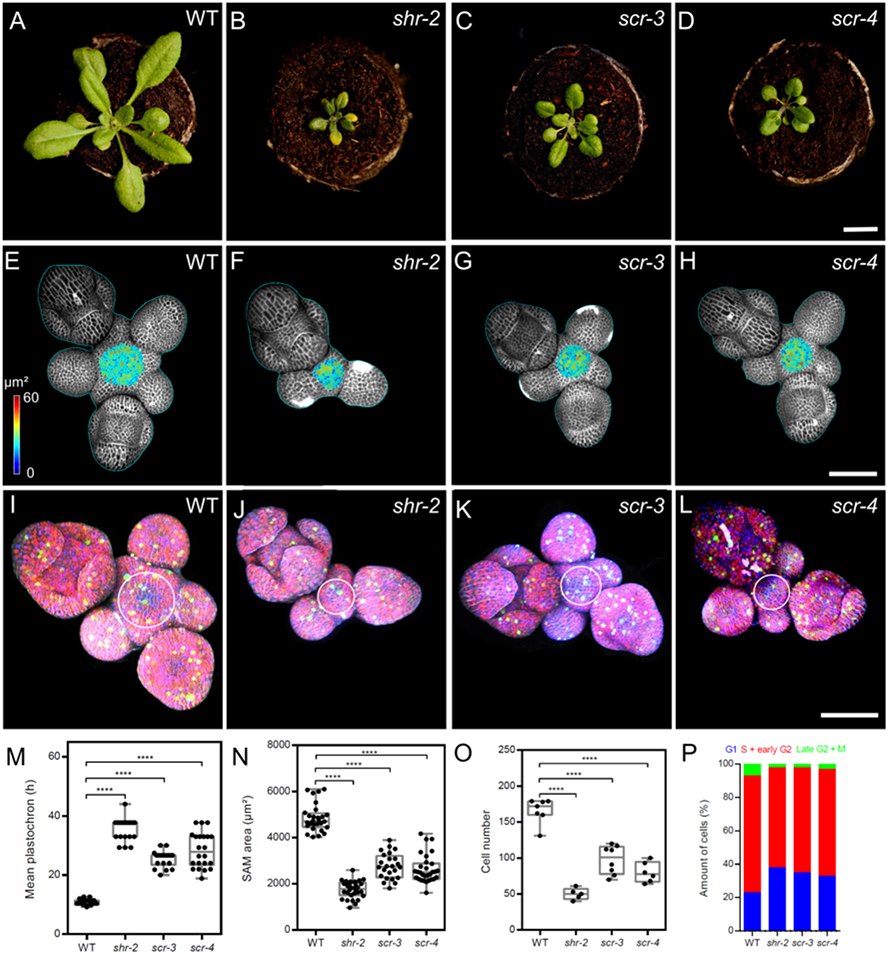
The *shr* and *scr* mutant phenotypes. **(A-D)** Top view of 21-day-old rosettes from WT (col-0) **(A)**, *shr-2* mutant **(B)**, *scr-3* mutant **(C)** and *scr-4* mutant **(D)**. Scale bar represents 1cm. **(E-H)** Heat-map quantification of the cell area in the meristem region at 5 weeks after germination from WT (n=7) **(E)**, *shr-2* mutant (n=5) **(F)**, *scr-3* mutant (n=8) **(G)** and *scr-4* mutant (n=6) **(G)**. Cell walls were stained with PI (gray). Scale bar represents 50 µm. **(I-L)** Representative 3D projection of shoot apical meristems at 5 weeks after germination expressing the three PlaCCI markers: *pCDT1a:CDT1a-eCFP* (blUE), *pHTR13:pHTR13-mCherry* (red), and *pCYCB1;1:NCYCB1;1-YFP* (green) in WT (n=11) **(I)**, *shr-2* mutant (n=4) **(J)**, *scr-3* mutant (n=7) **(K)** and *scr-4* mutant (n=6) **(L)**. White circles in **(I)**, **(J)**, **(K)** and **(L)** mark the meristem region. Cell walls were stained with DAPI (gray). Scale bar represents 50 µm. **(M)** Quantification of shoot apical meristem size at 5 weeks after germination from WT (n=28), *shr-2* mutant (n=30), *scr-3* mutant (n=24) and *scr-4* mutant (n=30). **(N)** Mean inflorescence plastochron in WT (n=21), *shr-2* mutant (n=20), *scr-3* mutant (n=18), and *scr-4* mutant (n=22). **(O)** Quantification of epidermal cell number in the meristem region of WT (n=11), *shr-2* mutant (n=4), *scr-3* mutant (n=7) and *scr-4* mutant (n=6). **(P)** Quantification of cells in different cell cycle phases in the meristem region (area surrounded by white circles in **(I)**, **(J)**, **(K)** and **(L)**) of WT (n=11), *shr-2* mutant (n=4), *scr-3* mutant (n=7) and *scr-4* mutant (n=6). Asterisks indicate a significant difference (^∗∗∗∗^p < 0.0001: Statistically significant differences were determined by Student’s *t*-test).

### SHR and SCR promote auxin signalling in the SAM

Since organ initiation sites are determined by auxin accumulation and signaling (Vernoux *et al*., 2011b), we asked if auxin accumulation is affected by loss of *SHR* or *SCR* functions, which would explain the observed delayed lateral organ initiation in *shr* and *scr* mutants (Fig. 1M). We introduced the auxin transcriptional output reporter *pDR5v2:3xYFP-N7* (Heisler et al., 2005) into the *shr-2* and *scr-4* mutants and assessed the number of domains with auxin output maxima compared to WT (Fig. 2A-C, arrowheads). We found that the average number of *DR5* positive domains in SAMs of *shr-2* and *scr-4* was significantly reduced (Fig. 2D), indicating that SHR and SCR functions are required to establish auxin maxima in the SAM.

**Fig. 2:**
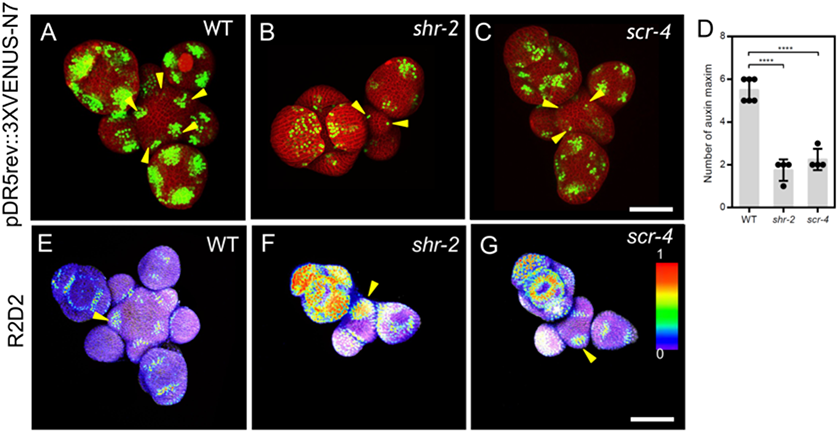
SHR and SCR functions modulate auxin signalling in the shoot apical meristem. **(A-C)** Representative 3D projection of shoot apical meristems at 5 weeks after germination expressing the auxin response reporter *pDR5rev:3XVENUS-N7* in WT (Col-0) (n=6) **(A)**, *shr-2* mutant (n=4) **(B)** and *scr-4* mutant (n=4) **(C)**. Yellow arrowheads in **(A)**, **(B)** and **(C)** show primordia with pDR5rev:3XVENUS-N7 expression. Cell walls were stained with PI (red). Scale bar represents 50 μm. **(D)** Quantification of auxin maxima in WT (n=6), *shr-2* mutant (n=4) and *scr-4* mutant (n=4). **(E-G)** Representative 3D projection of shoot apical meristems at 5 weeks after germination expressing the auxin input sensor R2D2 showing DII/mDII ratio intensity in WT (Col-0) (n=11) **(E)**, *shr-2* mutant (n=9) **(F)** and *scr-4* mutant (n=6) **(G)**. Yellow arrowheads in **(E)**, **(F)** and **(G)** show primordia with low auxin. Cell walls were stained with DAPI (gray). Scale bars represents 50 μm. Asterisks indicate a significant difference (^∗∗∗∗^p < 0.0001: Statistically significant differences were determined by Student’s *t*-test). Error bars display SD.

To closely evaluate how the distribution of auxin was affected in the SAM during the loss of SHR and SCR activities, we further evaluated auxin levels in the different meristem domains using the R2D2 auxin input sensor (Liao *et al*., 2015). R2D2 is a degron-based ratiometric auxin reporter consisting of an auxin-dependent degradation domain II (DII) of an Aux/IAA protein fused to Venus, and a mutant auxin nondegradable domain II (mDII) fused to tdTomato. The ratio of the fluorescence intensities between Venus and tdTomato represents a proxy for the level of auxin in every cell, with high ratios indicating low auxin levels and vice versa. We observed that the domains of low auxin in meristems of *shr-2* and *scr-4* mutants were expanded and sometimes extended beyond lateral organ boundaries into primordia (Fig. 2E-G, arrowheads). Altogether, these results show that SHR and SCR functions are required to maintain the proper distribution of auxin within the SAM. This safeguards the regular generation of auxin maxima in the peripheral zone, which is a prerequisite for organ initiation.

### SHR and SCR expression in shoot and flower meristems

To unravel the precise expression patterns of SHR and SCR in the meristem, we made use of previously established translational reporter lines which functionally complement the corresponding *shr* or *scr* mutants (Long et al., 2017). Using confocal microscopy, we imaged *pSHR:YFP-SHR* transgenic plant meristems and studied SHR expression in the inflorescence meristems at 36 DAG. YFP-SHR was detected in initiating flower primordia, in floral organ primordia and in diverse flower organs (Fig. 3A). The transcriptional reporter *pSHR:ntdTomato* (Möller et al., 2017) shows expression from P1 onwards in the third meristematic cell layer (L3) and, weakly, in the L2 of older primordia (Fig. 3C; Suppl Fig. 2A). No *SHR* expression was observed in the center of the SAM or floral meristems (Fig. 3A, left inset), or stem cell domains (Fig. 3A, right inset, asterisks). The translational reporter *pSHR:YFP-SHR* (Fig. 3A and D) revealed that SHR protein localized throughout L3 cells and appeared to be preferentially confined to the nuclei in L2 cells and also in L1 cells, where *SHR* is normally not expressed (Fig. 3C). Based on the combined observation of *SHR* transcriptional activity and the SHR protein localization, we conclude that SHR, similar to its activity in RAM, acts as a mobile transcription factor moving from the inner cell layers to the outermost cell layers of the organ primordia to mediate proper SAM development. We then evaluated the expression of *SCR* in the shoot meristem. A transcriptional reporter for *SCR* (*pSCR:H2B-YFP*) was previously described to be expressed in the QC and the root endodermis (Xu *et al*., 2006). In the SAM, we found fluorescent signal only in differentiated vasculature of the shoot, and in a patchy pattern in some flower primordia (Suppl Fig. 2B, inset). Since the transcriptional *SCR* reporter might lack control elements that contribute to the native *SCR* expression pattern, we used a translational reporter line, *pSCR:SCR-YFP* that was previously shown to complement all *scr* mutant phenotypes (Long et al., 2017). We observed SCR-YFP fluorescence in the nuclei of L1 cells in the central zone of the SAM and further extended into the deeper meristem layers specifically of the peripheral zone and lateral organ primordia (Fig. 3B, inset). Notably, SCR-YFP was lacking in the deeper regions of the meristem center and the rib meristem. In lateral organ primordia, *SCR* expression starts from the L1 and L2, and extends into deeper regions during further development. Stronger expression was found in the boundary region that separates lateral organ primordia from the remainder of the SAM (Fig. 3E). Thus, SHR and SCR proteins are found in partially overlapping domains during lateral organ primordia development. Importantly, *SHR* is missing from the SAM center, where *SCR* is expressed in the L1 (compare Fig. 3A, right inset, with Fig. 3B, inset).

**Fig. 3:**
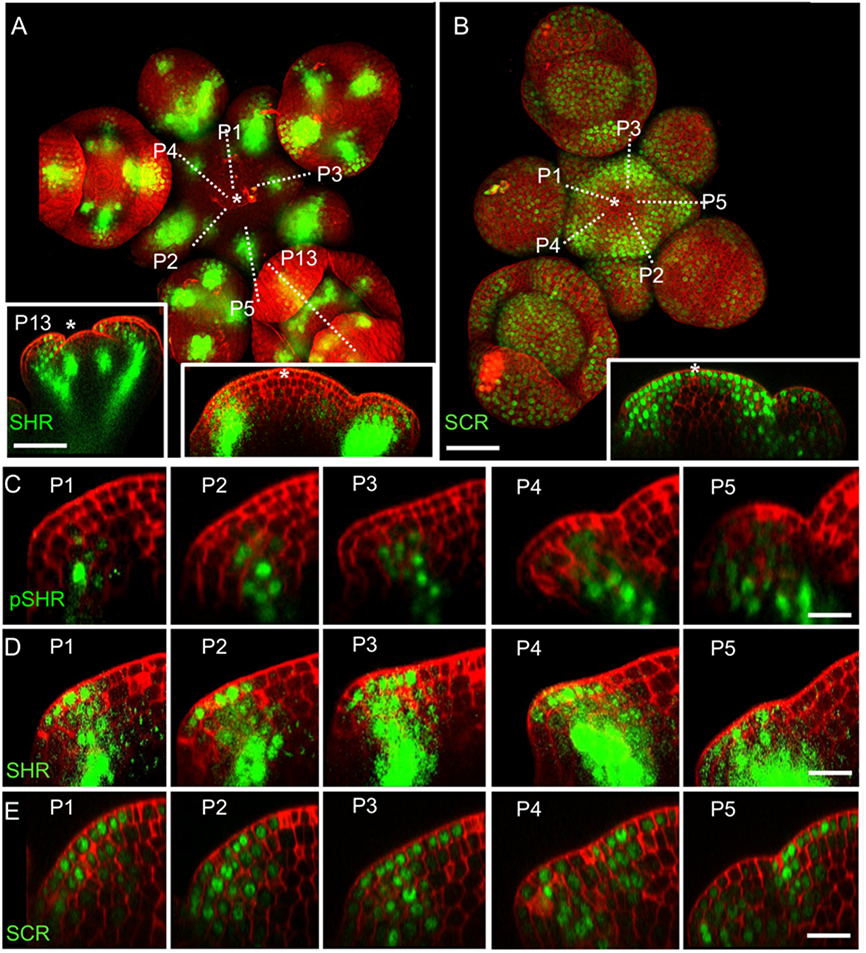
Expression patterns of SHR and SCR in the shoot apical meristem. **(A)** Representative 3D projection of SAM at 5 weeks after germination expressing *pSHR:YFP-SHR* reporter (green) (n=30). The lower right inset shows a longitudinal optical section through the middle of the SAM. The lower left inset shows a longitudinal optical section through the middle of primordia 13 (representative section orientation shown by dotted line). Cell walls were stained with DAPI (red). Scale bar represents 50 µm. **(B)** Representative 3D projection of SAM at 5 weeks after germination expressing *pSCR:SCR-YFP* reporter (green) (n=25). The lower right inset shows longitudinal optical section through the middle of the SAM. Cell walls were stained with PI (red). Scale bars represent 50 µm. **(C)** Longitudinal optical sections through the middle of five successive primordia expressing *pSHR:ntdTomato* reporter (green) (representative section orientation shown by dotted line in Suppl Fig. 2A). Cell walls were stained with DAPI (red). Scale bar represents 20 µm. **(D and E)** Longitudinal optical sections through the middle of five successive primordia expressing *pSHR:YFP-SHR* reporter (green) **(C)** and *pSCR:SCR-YFP* reporter (green) **(D)** (representative section orientation shown by dotted line in **(A)** and **(B)**, respectively). Scale bars represent 20 µm. P= Primordium.

### SHR and SCR form protein complexes in the SAM

SHR and SCR localize in an overlapping domain during lateral organ development (Fig. 3D and E), and SCR and SHR co-localized predominantly in nuclei of cells in the L1 and L3 of lateral organ primordia (Fig. 4A and C). Previous *in vivo* FRET-FLIM studies demonstrated direct interaction between SHR and SCR in the RAM (Long et al., 2017). We now investigated if SHR and SCR also interact in the SAM. For FRET-FLIM studies, we used YFP-SHR as fluorescent donor and SCR-RFP as acceptor. Reductions in the fluorescence lifetime of YFP due to resonance energy transfer to RFP are recorded as a quantitative readout for FRET, indicative of interaction between YFP-SHR and SCR-RFP. We used SAMs at 36 DAG for FRET measurements between YFP-SHR and SCR-RFP in lateral organ primordia, petal primordia and sepal primordia. As a negative control for non-interacting proteins, we used SAMs from plants coexpressing *pSCR:SCR-YFP* with *pSCR:SCR-RFP*, since SCR proteins do not homodimerize (Long et al., 2017) (Fig. 4B and D). The YFP-SHR donor-only sample showed a fluorescence lifetime of 2.75 ± 0.02ns (Fig. 4E), which significantly decreased by approximately 0.3 ± 0.03ns in cells where SCR-RFP was coexpressed (Fig. 4E), whereas samples with SCR-RFP and SCR-YFP coexpression showed no significant changes in mean fluorescence lifetime compared to the SCR-YFP donor-only control (Fig. 4F). These results show that the SCR and SHR proteins physically interact in the SAMs of Arabidopsis.

**Fig. 4:**
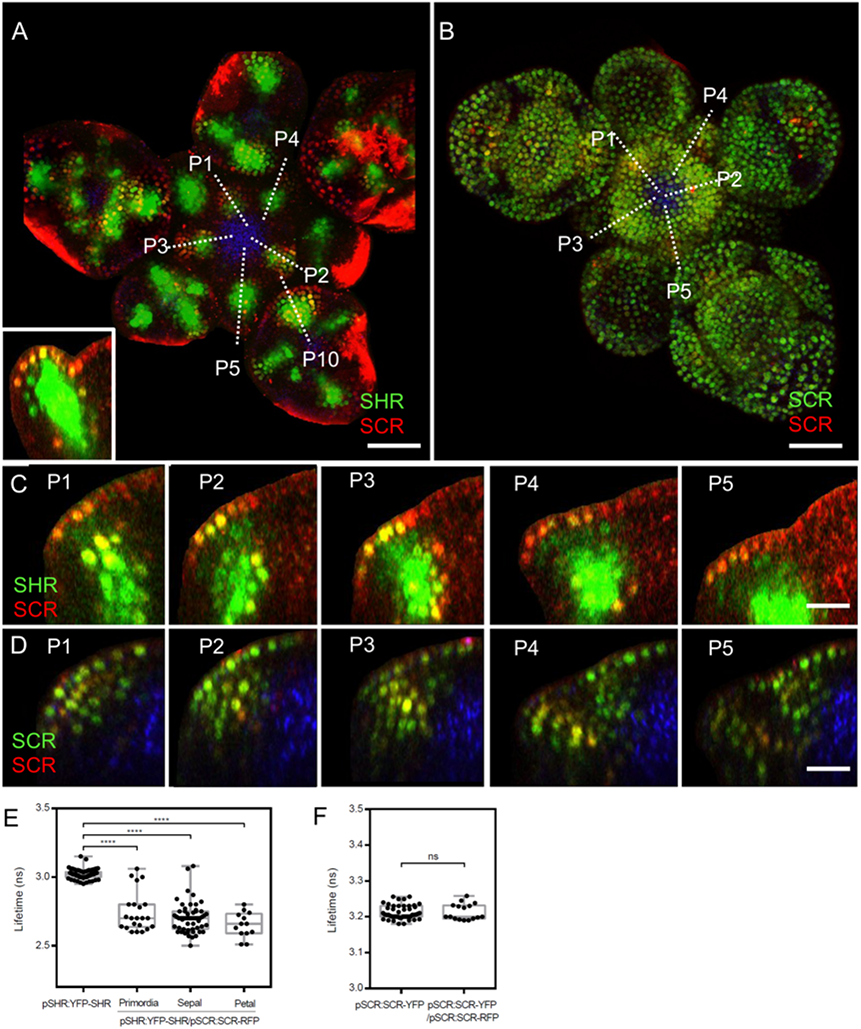
*In vivo* FRET–FLIM quantification of SHR-SCR association in the SAM. **(A-B)** Representative 3D projection of SAM at 5 weeks after germination coexpressing *pSHR:YFP-SHR* reporter (green) and *pSCR:SCR-RFP* reporter (red) (n=17) **(A)** (the lower left inset shows a longitudinal optical section through the middle of primordia 10; representative section orientation shown by dotted line) and *pSCR::SCR:YFP* reporter (green) and *pSCR::SCR:RFP* reporter (red) (n=11) **(B)**. Chlorophyll (blue). Scale bars represent 50 µm. **(C-D)** Longitudinal optical sections through the middle of five successive primordia coexpressing *pSHR:YFP-SHR* reporter (green) and *pSCR:SCR-RFP* reporter (red) **(C)**, and *pSCR::SCR:YFP* reporter (green) and *pSCR::SCR:RFP* reporter (red) **(D)** (representative section orientation shown by dotted line in **(A)** and **(B)** respectively). Chlorophyll (blue). Scale bars represent 20 µm. **(E)** Average lifetime of YFP-SHR when expressed alone (pSHR:YFP-SHR (*n* =75)), or coexpressed together with SCR-RFP (pSHR:YFP-SHR/pSCR:SCR-RFP) in lateral organ primordia (n=22), sepal primordia (n=55) and petal primordia (n=13). **(F)** Average lifetime of SCR-YFP when expressed alone (pSCR:SCR-YFP), or coexpressed together with SCR-RFP (pSCR:SCR-YFP/pSCR:SCR-RFP). Asterisks indicate a significant difference (^∗∗∗∗^p < 0.0001: Statistically significant differences were determined by Student’s *t*-test, ns= no significant difference). P= primordium.

### A SHR-SCR heteromer regulates expression of *SCR* in lateral organ primordia

We next tested if SHR regulates *SCR* expression. We compared the SCR protein localization using the *pSCR:SCR-YFP* reporter line in WT and the *shr-2* mutant background. SCR protein levels remained unaffected in the central zone of the SAM, where *SHR* is normally not expressed (Fig. 5B, C and G). However, in the lateral organ primordia, SCR protein level was strongly reduced in *shr-2* compared to WT (Fig. 5D, E and H). Analysis of *SCR* transcript levels by qRT-PCR corroborated a significant decrease in *shr-2* mutants shoot meristems (Fig. 5I). We conclude that SHR acts as a transcriptional activator of *SCR* in the SAM, but that *SCR* expression in the meristem center is SHR-independent. Importantly, the expression pattern of *pSHR:ntdTOMATO* does not entirely overlap with the expression pattern of *pSCR:SCR-YFP*, indicating that SHR acts as a mobile protein also in the SAM (Suppl Fig. 6A,D). We then tested if SCR restricts movement of SHR protein by nuclear retention in the SAM or lateral organ primordia. Analysis of SAMs coexpressing *pSHR:SHR-YFP* and *pSCR:SCR-RFP* showed that SHR-YFP is enriched in the nuclei of cells coexpressing SCR-RFP at the lateral organ primordia (Fig. 5A inset and F). This enrichment could indicate that SCR entraps SHR in the nuclei, thus restricting its further intercellular movement.

**Fig. 5:**
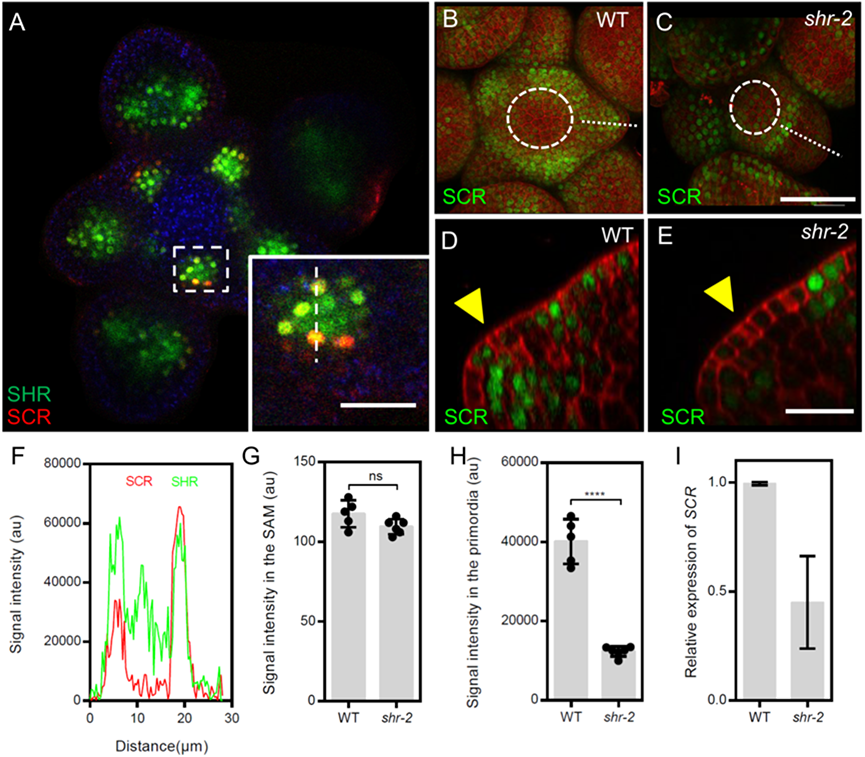
SHR regulates *SCR* expression in the SAM. **(A)** Representative transversal confocal image of SAM at 5 weeks after germination coexpressing the *pSHR:SHR-YFP* reporter (green) and the *pSCR:SCR-RFP* reporter (red) (n=5). Chlorophyll (blue). The lower right inset shows an optical section view with high magnification of the area surrounded by white dashed rectangle in **(A)**. Scale bar represents 50 µm. **(B and C)** Representative 3D projection of SAM at 5 weeks after germination expressing the *pSCR:SCR-YFP* reporter (green) in WT (n=16) **(B)** and *shr-2* mutant (n=10) **(C)**. Cell walls were stained with PI (red). Scale bar represents 50 µm. **(D and E)** Longitudinal optical sections through the middle of primordia shown by dotted line in **(B)** and **(C)** respectively. Yellow arrowheads in **(D)** and **(E)** show primordia with SCR-YFP expression. Scale bar represents 20 µm. **(F)** Intensity plot profile of SHR-YFP (green) signal and the SCR-RFP (red) signal in the area crossed by white dashed line in **(A)** inset. **(G and H)** Quantification of SCR-YPF signal intensity in the meristem center (area surrounded by white dashed circles in **(B)** and **(C)**) **(G)** and in the primordia **(H)**. **(I)** Quantitative real-time reverse transcriptase PCR analysis showing the relative expression levels of *SCR* in WT and *shr-2* mutant SAMs. The expression level in Col-0 is set to 1. Expression levels were normalized using AT4G34270 and AT2G28390. Asterisks indicate a significant difference (^∗∗∗∗^p < 0.0001: Statistically significant differences were determined by Student’s *t*-test, ns= no significant difference). Error bars display SD.

### Roles of *JKD* in the SAM

The BIRD-family transcription factor JACKDAW (JKD) can interact with SHR and SCR and form multiprotein complexes (Long et al., 2015b, Long et al., 2017, Welch *et al*., 2007), here we investigated if JKD also affects SAM development. SAM sizes of *jkd-4* mutants were increased compared to WT, due to a significant rise in cell number (Fig. 6A-D, and Suppl Fig. 3A). For expression studies, we used the transcriptional reporter *pJKD:YFP-RFP* and the translational *pJKD:JKD-YFP* reporter that has been previously shown to complement the *jkd-4* mutant phenotype (Long et al., 2017). *JKD* was found to be expressed only in some cells of the peripheral zone of the SAM, and in the abaxial regions of flower primordia (Fig. 6E-F and I). The expression patterns of both transcriptional and translational reporter lines were mostly identical (Fig. 6E-F), indicating that *JKD* RNA and protein are found in the same cells, and that JKD protein is not mobile. The JKD expression domain expands from few cells at the abaxial side of P1 into a ring-like abaxial domain in sepals, and at later stages, in stamen and carpel primordia (Fig. 6G and I). Based on the observed JKD expression (Fig 6E-F and I), we asked if JKD localization directly correlates with early sepal development. To do this, we used the strong *lfy-12* mutant where flower meristems fail to produce proper flower organs but instead generate mostly sepal-like organs (Weigel et al., 1992), and the *clv3-9* mutant, which has an excessively enlarged SAM and flowers with additional organs (Schlegel *et al*., 2021). JKD was confined to sepal primordia of *clv3-9* mutants and to the sepal-like organs of *lfy-12* mutants like in WT (Suppl Fig. 3C-D). We tested for potential molecular interactions of JKD with SCR in the SAM *in vivo* using FRET-FLIM. SAMs coexpressing *pJKD:JKD-YFP* and *pSCR:SCR-RFP* (Fig. 6J) showed significant fluorescence lifetime reductions of up to 0.17 ± 0.02ns, compared to the JKD-YFP donor-only which exhibited a lifetime of 3.02 ± 0.03ns (Fig. 6H). We conclude that SHR-SCR can form multimeric complexes with JKD in the SAM periphery, in floral organ primordia and in floral organs. JKD could modulate or even dampen activities of the SHR-SCR complexes, since JKD and SHR-SCR exert opposite effects on meristem size.

**Fig. 6:**
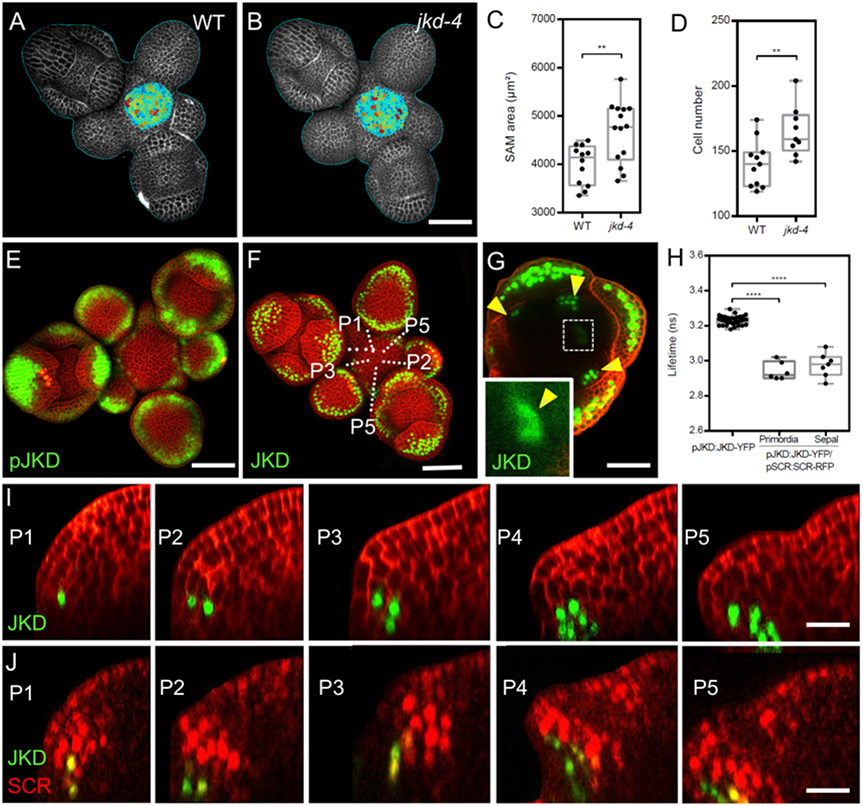
JKD functions and expression pattern in the SAM. **(A and B)** Heat-map quantification of the cell area in the meristem region at 5 weeks after germination from WT **(A)** (n=10) and *jkd-4* mutant **(B)** (n=10). Cell walls were stained with PI (gray). Scale bar represents 50 µm. **(C)** Quantification of SAM size at 34 days after germination (DAG) from WT (n=12) and *jkd-4* mutant (n=14). **(D)** Quantification of epidermal cell number in the meristem region of WT **(A)** (n=10) and *jkd-4* mutant **(B)** (n=10). **(E and F)** Representative 3D projection of SAM at 5 weeks after germination expressing *pJKD:YFP-RFP* reporter (green) (n=9) **(E)** and *pJKD:JKD-YFP* reporter (green) (n=24) **(F)**. Cell walls were stained with PI (red). Scale bars represent 50 µm. **(G)** Transversal optical through the primordium 16 from an inflorescence apex. Yellow arrowheads show petal primordia with *pJKD:JKD-YFP* expression (green). The lower left inset shows an optical view with high magnification of the area surrounded by a white dashed rectangle. Cell walls were stained with PI (red). **(H)** Average lifetime of JKD-YFP when expressed alone (*pJKD:JKD-YFP* (n =38)), or coexpressed together with SCR-RFP (*pJKD:JKD-YFP/pSCR:SCR-RFP*) in lateral organ primordia (n=6) and sepal primordia (n=7). **(I)** Longitudinal optical sections through the middle of five successive primordia (representative section orientation shown by dotted line in **(A)**) expressing *pJKD:JKD-YFP* reporter (green). Cell walls were stained with PI (red). Scale bar represents 20 µm. **(J)** Longitudinal optical sections through the middle of five successive primordia (representative section orientation shown by dotted line in Suppl Fig. 4F) coexpressing *pJKD:JKD-YFP* reporter (green) and *pSCR:SCR-RFP* reporter (red) (n=7). Scale bar represents 20 µm. Asterisks indicate a significant difference (^∗∗∗∗^p < 0.0001; ^∗∗^p < 0.001: Statistically significant differences were determined by Student’s *t*-test). P= primordium.

### The SHR-SCR-JKD complex triggers cell division in lateral organ primordia by regulating *CYCD6;1* expression

Lateral organ initiation in the SAM periphery is controlled by *CYCLIN* genes, which are key regulators of cell division (Dewitte *et al*., 2003). We examined the expression patterns of *pCYCD1;1:GFP, pCYCD2;1:GFP, pCYCD3;2:GFP, pCYCD3;3:GFP, pCYCD5;1:GFP, pCYCD6;1:GFP, pCYCD7;1:GFP* and *pCYCB1;1:CYCB1;1-GFP (Ubeda-Tomás et al., 2009).* Amongst these, only *CYCD1;1*, *CYCD3;2*, *CYCD3;3*, *CYCD6;1* and *CYCB1;1* were visibly expressed in the SAM (Fig. 7A; Suppl Fig. 4A-D and G). *CYCD1;1*, *CYCD3;2* and *CYCB1;1* exhibited a patchy expression pattern in sepal primordia (Suppl Fig. 4A-B and D), while *CYCD3;3* was more uniformly distributed with an enrichment in lateral organ primordia (Suppl Fig. 4C). *pCYCD6;1:GFP* expression was detected in lateral organ primordia and floral organs in sepal primordia from all stages, but not at the center of the SAM (Fig. 7A,C,E,L and O). Notably, *pCYCD6;1:GFP* was predominantly expressed in the region where expression of SHR, SCR and JKD overlapped (Fig. 7J-O; Suppl Fig. 4E-S). We tested if expression of *pCYCD6;1:GFP* is affected when *SHR* function is compromised, and observed a strong reduction in *pCYCD6;1:GFP* expression level in the sepal primordia of *shr-2* and its complete absence from lateral organ primordia in the *shr-2* mutant, while *pCYCD6;1:GFP* was detected in the L3 of WT lateral organ primordia (Fig. 7A-F). In the root meristem, *CYCD6;1* specifically controls periclinal cell divisions and the formation of new tissue layers (Long et al., 2015b). We found that the number of cell layers in the L3 of lateral organ primordia is strongly reduced in *shr-2* mutants compared to WT (Fig. 7G-I), consistent with the SHR-SCR-JKD complex acting through *CYCD6* to inducte periclinal cell divisions in the SAM.

**Fig. 7:**
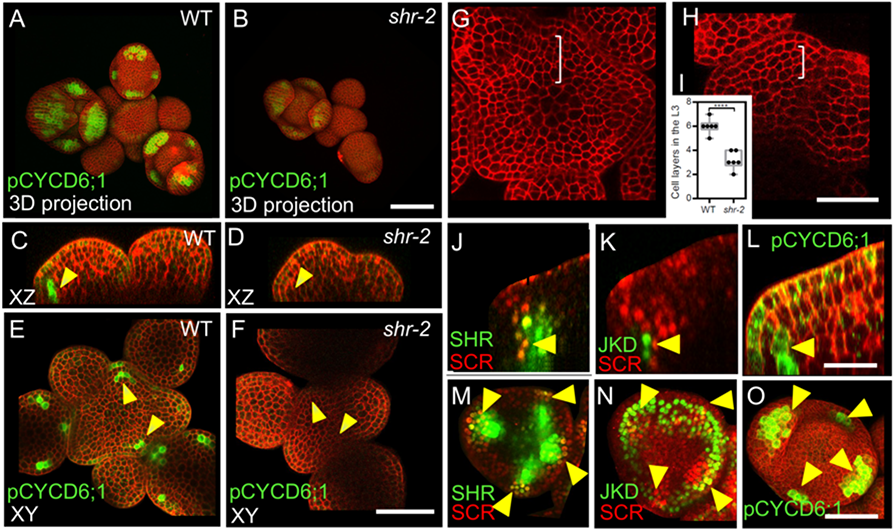
SHR regulates *CYCD6;1* expression in the SAM. **(A and B)** Representative 3D projection of SAM at 5 weeks after germination expressing *pCYCD6;1:GFP* reporter (green) in WT (n=11) **(A)** and *shr-2* mutant (n=6) **(B)**. Cell walls were stained with PI (red). Scale bar represents 50 µm. **(C and D)** Longitudinal optical sections of **(A)** and **(B)** respectively. **(E and F)** Transversal optical sections of **(A)** and **(B)** respectively. Scale bar represents 50 µm. **(G and H)** Transversal optical sections of the inflorescence apex at 5 weeks after germination from WT **(G)** and *shr-2* mutant **(H)**. **(I)** Quantitative comparison of cell files within the L3 in optical sections of the inflorescence apex from WT (n=6) and *shr-2* mutant (n=6). **(J-L)** lateral organ primordia showing coexpression of *pSHR:SHR-YFP* reporter (green) and *pSCR:SCR-RFP* reporter (red) **(J)**, *pJKD:JKD-YFP* reporter (green) and *pSCR:SCR-RFP* reporter (red) **(K)** and *pCYCD6;1:GFP* reporter (green) **(L)**. Scale bar represents 20 µm. **(M-O)** Florescence meristem stage 4 of flower development showing coexpression of *pSHR:SHR-YFP* reporter (green) and *pSCR::SCR-RFP* reporter (red) **(M)**, *pJKD:JKD-YFP* reporter (green) and *pSCR::SCR-RFP* reporter (red) **(N)** and *pCYCD6;1:GFP* reporter (green) **(O)**. Scale bar represents 20 µm. Asterisks indicate a significant difference (^∗∗∗∗^p < 0.0001; analyzed by Student’s t test).

### Auxin induces *SHR* and *SCR* expression in the SAM

Since we observed that *SHR* and *SCR* control SAM size and lateral organ initiation which is triggered by local auxin accumulation, we investigated SHR-SCR interaction with auxin distribution and/or signaling. We wanted to first examine if *SHR* and *SCR* expression patterns correlated with that of auxin distribution using the functional *pPIN1:PIN1-GFP* reporter (Heisler et al., 2005). PIN1 is an auxin efflux transporter which controls the flow and the distribution of auxin across the SAM (Benková et al., 2003) (Suppl Fig. 6C and F). The expression patterns of *SHR* and *SCR* in primordia strongly overlapped with that of *PIN1* during primordia initiation and development (Fig. 3A-D; Suppl Fig. 6A-F). This indicates that *SHR* and *SCR* might act downstream in response to auxin signaling. *In silico* analysis of the *SHR* promoter sequence revealed the presence of two putative auxin-response element (AuxRE) core motifs (TGTCTC) (Fig. 10A). These motives are thought to be sufficient for recruitment of ARF transcription factors (Schlereth *et al*., 2010, Zhao *et al*., 2010, Ulmasov *et al*., 1999), which are the downstream TFs released after auxin dependent degradation of IAAs (Paque & Weijers, 2016). This suggests that *SHR* might be induced by ARF upon auxin signaling. We then tested if auxin regulates expression of *SHR* and *SCR* in the SAM and treated WT inflorescences with 10µM indole-3-acetic acid (IAA). Within 12h of hormone application, the expression levels of both *SHR* and *SCR* increased to about 300% of untreated control levels. *SHR* expression increased slightly earlier than that of *SCR*, further supporting our previous observation that *SHR* activates *SCR* expression (Fig. 8A). To uncover the spatial distribution of *SHR* and *SCR* in response to exogenous auxin application, we analyzed SAMs expressing *pSHR:SHR-YFP* and *pSCR:SCR-YFP* after treatment with 10µM of the synthetic auxin analogue 2,4D, a form of auxin that can diffuse into cells. We found that expression visibly increased within 2-5h after the start of the hormone treatment (Fig. 8B-G). Importantly, the expression patterns of both *SHR* and *SCR* were not strongly altered. Since we observed an overlap of auxin accumulation domains with those of *SHR* and *SCR,* and also established that auxin induces *SHR* and *SCR* expression, we tested if changes in auxin transport and local distribution in turn affected *SHR* and *SCR* expression patterns. We inhibited polar auxin transport and hence its distribution in tissues using N-1-naphthylphthalamic acid (NPA). The mock control SAMs expressing *pPIN1:PIN1-GFP* showed a wide distribution of *PIN1-GFP*, which relocated into a ring shaped domain in deeper regions of the meristem 3d after treatment with 100µM NPA (Fig. 8P-S). Concomitantly, the expression pattern of *SHR-YFP* changed from being associated with primordia into a ring shaped domain, reflecting the rearrangement of the *PIN1* expression domain (Fig. 8H-K). Similar changes in expression pattern were found for *SCR* upon NPA treatment (Fig. 8L-O). In addition, NPA treatment resulted in enlarged expression domains for *pCYCD6;1:GFP*, a key target gene of the SHR-SCR-JKD complex (Suppl Fig. 5A-D). Based on all these observations, we conclude that, in the SAM, *SHR* and *SCR* levels and expression pattern are to a large extent coordinated by auxin levels and distribution.

**Fig. 8:**
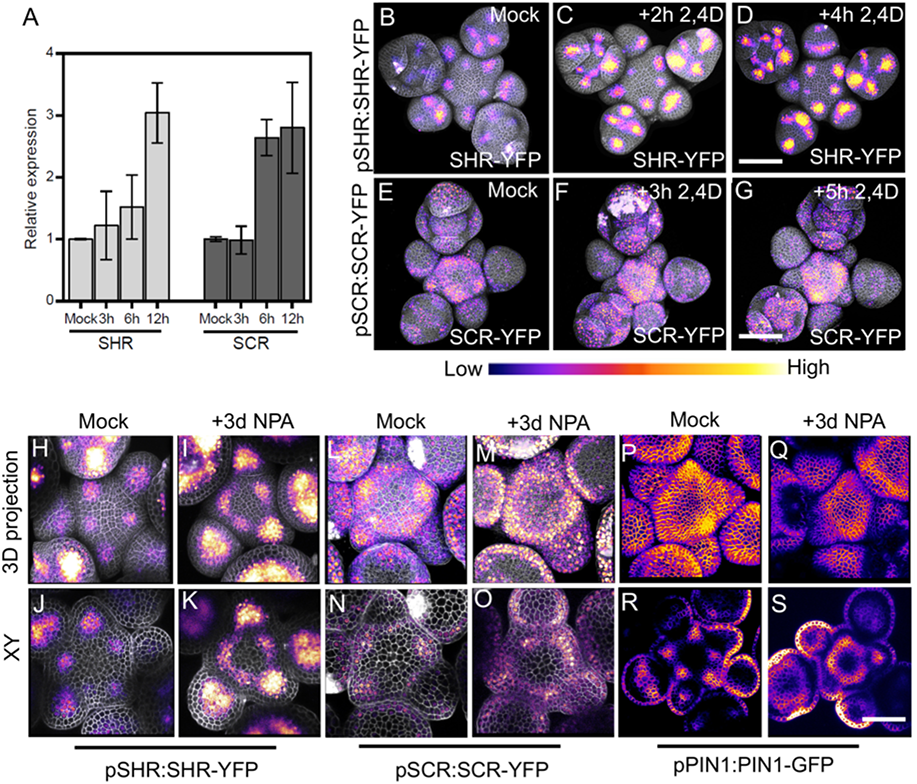
SHR and SCR expressions respond to auxin. **(A)** Quantitative real-time reverse transcriptase PCR analysis showing the relative expression levels of *SHR* and *SCR* expression in response to auxin (10 µm IAA) in WT SAMs. The expression level in Col-0 is set to 1 and error bars show standard deviation. Expression levels were normalized using AT4G34270 and AT2G28390. **(B-D)** Representative 3D projection of SAMs at 5 weeks after germination expressing *pSHR:SHR-YFP* reporter (magenta) in mock **(B)** (n=18), or after 2h (n=12) **(C)** or 4h 10µM 2,4D treatment (n=8) **(D)**. Fluorescence intensities were coded blue to yellow corresponding to increasing intensity. Scale bar represents 50 µm. **(E-G)** Representative 3D projection of SAMs at 5 weeks after germination expressing *pSCR:SCR-YFP* reporter (magenta) mock **(E)** (n=11), after 3h (n=6) **(F)** or after 5h 10µM 2,4D treatment (n=7) **(G)**. Fluorescence intensities were coded blue to yellow corresponding to increasing intensity. Scale bar represents 50 µm. **(H and I)** Representative 3D projection of SAMs at 5 weeks after germination expressing *pSHR:SHR-YFP* reporter (magenta) three days after mock (n≥40) **(H)** or 100µM NPA treatment (n≥60) **(I)**. **(J and K)** Transversal optical sections of **(H)** and **(I)** respectively. **(L and M)** Representative 3D projection of SAMs at 5 weeks after germination expressing *pSCR:SCR-YFP* reporter (magenta) three days after mock (n=5) **(L)** or 100µM NPA treatment (n=10) **(M)**. **(N and O)** Transversal optical sections of **(L)** and **(M)** respectively. **(P and Q)** Representative 3D projection of SAMs at 5 weeks after germination expressing *pPIN1:PIN1-GFP* reporter (magenta) three days after mock (n=9) **(P)** or 100µM NPA treatment (n=16) **(Q)**. **(R and S)** Transversal optical sections of **(P)** and **(Q)** respectively.

### MONOPTEROS regulates *SHR* and *SCR* expression in the SAM

We have shown that auxin regulates *SHR* expression likely through ARFs (Fig 8A). The ARF MONOPTEROS (MP) is required to promote expression of *SHR* and *SCR* in the early embryo (Möller et al., 2017). In the SAM, the expression patterns of *MP, SCR* and *SHR* partially overlap in the primordia (Fig. 3D and E; Suppl Fig. 6A-B and D-E). We then analyzed the expression of *pSHR:ntdTomato* in *mp-B4149,* a loss-of-function mutant for *MP*. In WT vegetative meristems at 7 DAG, *SHR* was expressed in the deeper region of the young leaf primordia, where the vasculature will eventually develop. *SHR* was expressed in a similar pattern in the *mp-B4149* mutants, albeit at lower levels (Fig. 9A-B), indicating that *MP* is required for normal *SHR* expression levels post embryogenesis. Since *mp-B4149* mutants fail to develop beyond the early seedling stage, we used the hypomorphic allele *mp-S319* to study later stages of development. *mp-S319* mutants display weaker phenotypes than *mpB-4149* null mutants and initiate an inflorescence meristem with, occasionally, some flower primordia. We used the transcriptional reporter *pSHR:ntdTOMATO* and the translational reporter *pSHR:SHR-YFP* to distinguish between effects of auxin on *SHR* promoter activity and posttranscriptional effects leading to altered protein localization. Compared to WT, *pSHR:ntdTOMATO* expression was strongly reduced in *mp-S319* inflorescences (Fig. 9C-F). Similar results were observed for the translational reporter *pSHR:SHR-YFP*, and weak expression in a ring-shaped domain was found in cross sections through the inflorescence stem (Fig. 9K-N). Similarly, using *pSCR:SCR-RFP*, we observed reduced expression of *SCR* in the meristem periphery in *mp-S319*, which now overall resembled the *SHR* expression pattern (Fig. 9G-J). We therefore conclude that *MP* is required for maintaining normal expression of *SHR* and *SCR* expression in the meristem periphery and primordia. We then tested whether *MP* could promote *SHR* expression throughout the meristem, using an inducible MP-GR fusion protein expressed from the *Ubiquitin10* promoter (*pUBQ10:MP-GR*). Treatment of these plants with dexamethasone (DEX) allows the nuclear entry of MP-GR. Within 4 h of DEX treatment, ubiquitously expressed MP resulted in increased expression levels of *pSHR:SHR-YFP* and *pSHR:ntdTOMATO* while retaining its WT expression domain (Fig. 9O-P; Suppl Fig. 10Q and R), indicating that MP can increase *SHR* expression levels, but is not sufficient to control the precise expression pattern in the absence of additional auxin. Thus, MP exerts a quantitative regulatory control over *SHR* expression. To better understand the distinct roles of *SHR* and *MP* in organ initiation, we generated double mutants of *mp-S319* and *shr-2*. Unlike *mp-S319* that forms only fewer flower primordia than WT, the phenotype of the double mutant *mp-S319; shr-2* was more severe with naked inflorescences that lacked all organ primordia (Suppl Fig. 7A-D’’). We conclude that MP acts through SHR to promote lateral organ initiation.

**Fig. 9:**
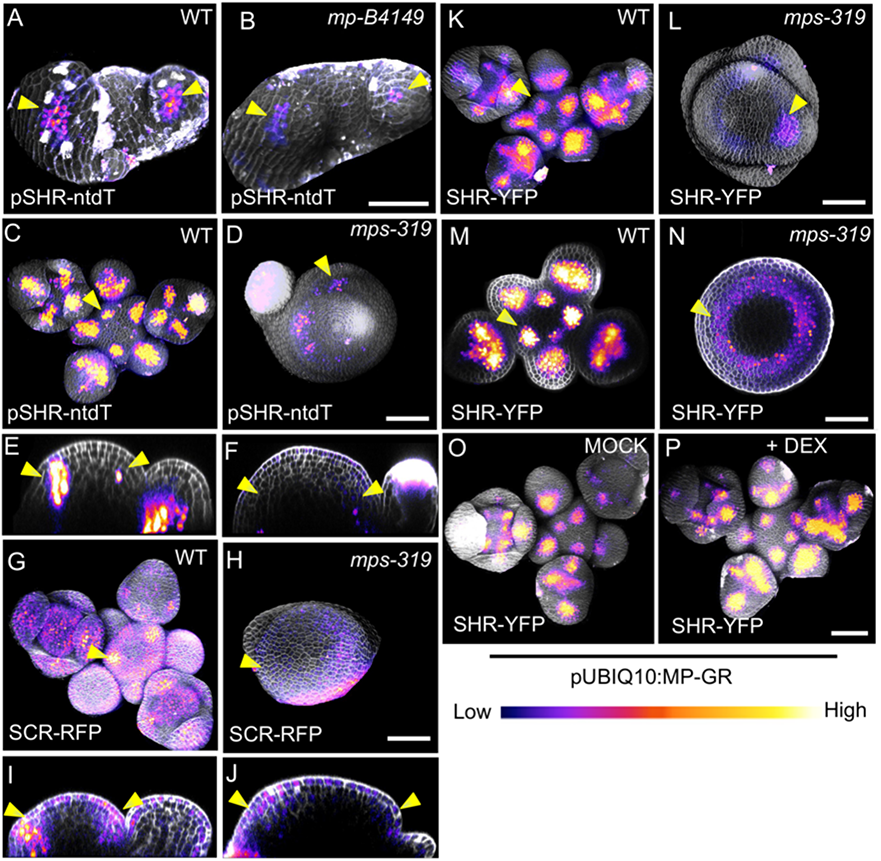
SHR and SCR act downstream of MP in the SAM. **(A and B)** Representative 3D projection of SAMs at 5 days after germination expressing *pSHR:ntdTomato* reporter (magenta) in WT (n=8) **(A)** and *mp-B4149* (n=5) **(B)** mutant vegetative meristems. Yellow arrowheads in **(A)** and **(B)** indicate the region where flower primordia initiate and *pSHR:ntdTomato* expression. Scale bar represents 50 µm. **(C and D)** Representative 3D projection of SAMs at 5 weeks after germination expressing *pSHR:ntdTomato* reporter (magenta) in WT (n=10) **(C)** and *mps-319* mutant (n=15) **(D)**. **(E and F)** Longitudinal optical section of **(C)** and **(D)** respectively. Yellow arrowheads in **(C)**, **(D)**, **(E)** and **(F)** indicate the region where flower primordia initiate and *pSHR:ntdTomato* expression. Scale bar represents 50 µm. **(G and H)** Representative 3D projection of SAMs at 5 weeks after germination expressing *pSCR:SCR-RFP* reporter (magenta) in WT (n=12) **(G)** and *mps-319* mutant (n=4) **(H)**. **(I and J)** Longitudinal optical section of **(G)** and **(H)** respectively. Yellow arrowheads in **(G)**, **(H)**; **(I)** and **(J)** indicate the region where flower primordia initiate and *pSCR:SCR-RFP* expression. Scale bar represents 50 µm. **(K and L)** Representative 3D projection of SAMs at 5 weeks after germination expressing *pSHR:SHR-YFP* reporter (magenta) in WT (n=9) **(K)** and *mps-319* mutant (n=6) **(L)**. **(M and N)** Transversal optical section of **(K)** and **(L)** respectively. Yellow arrowheads in **(K)**, **(L)**; **(M)** and **(N)** indicate the region where flower primordia initiate and *pSHR:SHR-YFP* expression. Scale bar represents 50 µm. **(O and P)** Representative 3D projection of shoot apical meristems at 5 weeks after germination expressing *pSHR:SHR-YFP* reporter (magenta) in *pUBIQ10:MP-GR* plants 4hr after mock (n=5) **(O)** or 4hr after DEX treatment (n=7) **(P)**. Fluorescence intensities were coded blue to yellow corresponding to increasing intensity. Scale bar represents 50 µm.

### AuxRE motifs in the *SHR* promoter control its activity

We showed that *SHR* expression is induced by the ARF MP (Fig. 9A-F and K-P; Suppl Fig. 10Q and R). To test if MP can directly activate *SHR* expression, we analyzed two candidate AuxREs (AuxRE1 and AuxRE2) in the *SHR* promoter. We tested the ability of the *SHR* promoter, with or without these AuxREs, to control the expression of the reporters mVenus or Luciferase upon Agrobacterium-mediated transient expression in *Nicotiana benthamiana* leaves. mVenus under control of the WT *SHR* promoter was co-infiltrated with *p35S:MP*, or *p35S:GUS* as a negative control. We observed strong *mVenus* expression from the WT *SHR* promoter in the presence of *p35S:MP*, which was about 6 fold higher than upon coexpression of *p35S:GUS* (Suppl Fig. 8B). Mutating AuxRE1 from “GAGACA” to “GAGCCA” (mAuxRE1) or “GTGCTC” (mAuxRE1-2) and AuxRE2 from “TGTCTC” to “TGGCTC“ (mAuxRE2) or “TGGAGA” (mAuxRE2-2) by site directed mutagenesis drastically reduced the response to *p35S:MP* coexpression, and mutating both AuxREs had an additive effect, resulting in no significant response to overexpression of *MP* (Suppl Fig. 8A-J and Suppl Fig. 9A-J). Similar results were obtained using a Luciferase assay system (Suppl Fig. 10A-H), indicating that MP interacts with these two AuxRE elements in the *SHR* promoter to regulate *SHR* expression. To investigate if these elements also contribute to *SHR* expression in Arabidopsis, we generated transgenic plants using either the WT *SHR* promoter, or the mutant versions carrying base changes in one or both AuxREs. At least ten independent transgenic lines were analyzed for each promoter construct. Expression of the WT *pSHR:mVenus-NLS* produced a strong expression in lateral organ primordia and floral meristems. However, mutations in AuxRE1 (mAuxRE1 or mAuxRE1-2) or AuxRE2 (mAuxRE2 or mAuxRE2-2) led to decreased mVenus expression in all organ primordia stages and the floral meristem (Fig. 10B-D; Suppl Fig 9K-M). When both AuxREs (mAuxRE1+2 or mAuxRE1-2+2-2) were mutated, mVenus expression was barely detectable (Fig. 10E; Suppl Fig 9N). We concluded that both AuxRE motifs function in an additive manner during ARF-dependent activation of *SHR* expression in the SAM. Notably, consistent with the expression in the SAM, mutations in AuxREs motifs in the *SHR* promoter also led to a decreased *mVenus* expression in the RAM compared to the WT *pSHR:mVenus-NLS,* where *SHR* and *MP* expression overlap (Suppl Fig. 11A-H).

**Fig. 10:**
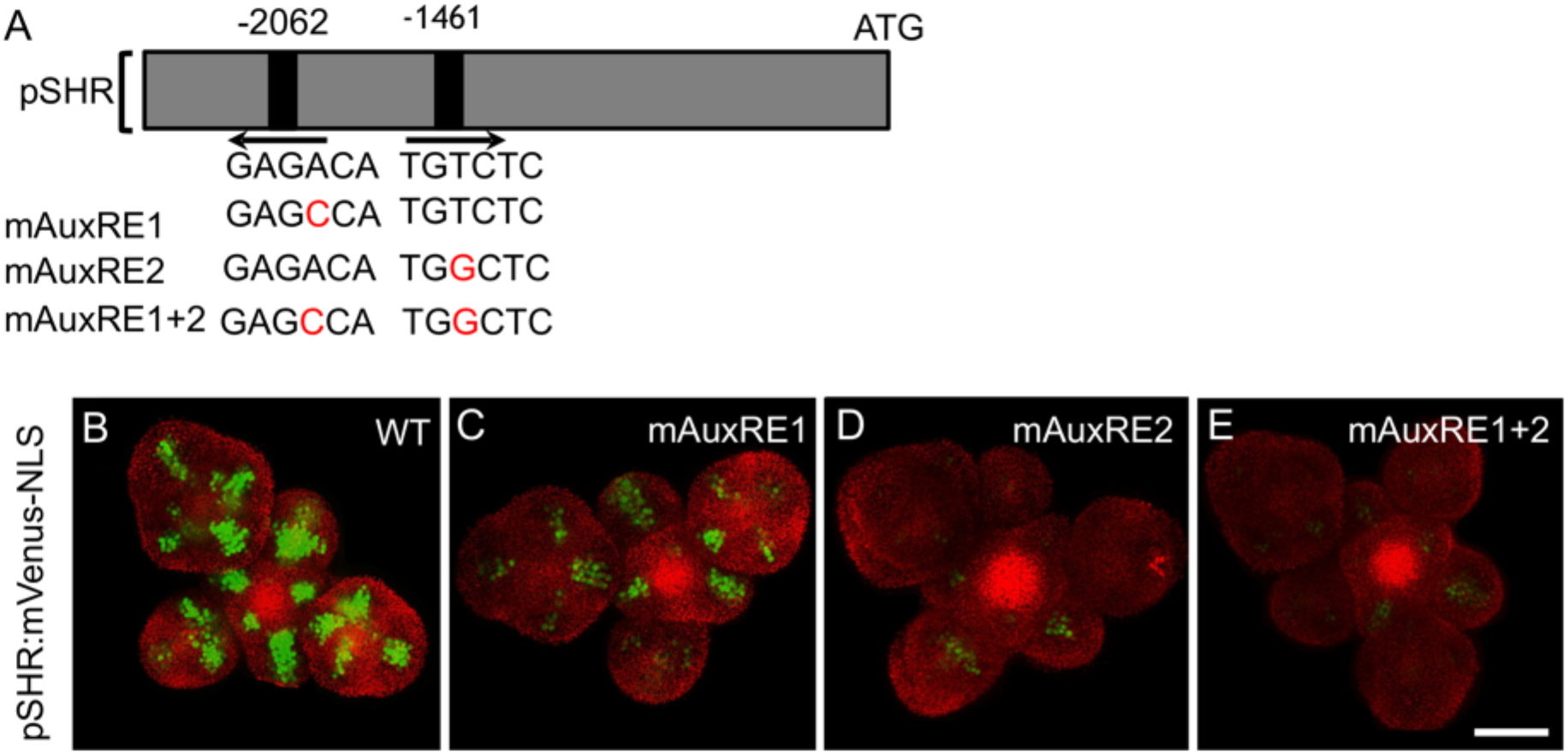
MP Regulates *SHR* expression in the SAM. **(A)** Schematic representation of the *SHR* promoter. The positions of two auxin response elements are shown. Overview of mutated promoter versions of *pSHR*. AuxREs were mutated and multiple combinations of these mutated motifs were combined into a single promoter. The original AuxRE sequence GAGACA was mutated to GAGCCA (mAuxRE1), the original AuxRE sequence TGTCTC was mutated to TGGCTC (mAuxRE2). **(B–E)** Representative 3D projection of SAMs at 5 weeks after germination expressing mVenus-NLS under the control of the wild-type *SHR* promoter (n=12) **(B)**, and under the control of the *SHR* promoter with mutations in AuxRE motifs mAuxRE1 (n=7) **(C)** mAuxRE2 (n=5) **(D)** and mAuxRE1+2 (n=11) **(E)**; Chlorophyll (red). Scale bar represents 50 µm.

### *SHR* regulates *LFY* expression in the SAM

It has been previously shown that MP directly activates *LFY* expression to confer floral identity to organ primordia (Yamaguchi et al., 2013). Since MP induces also *SHR* expression, we asked how *SHR* and *LFY* activities are coordinated. In WT, *SHR* and *SCR* were expressed two plastochrons prior to *LFY* in the primordia (Suppl Fig. 12A), and we hypothesized that *SHR* might mediate MP-dependent *LFY* expression. We first tested whether *LFY* expression was altered in *shr-2* mutants. mRNA quantification via qRT-PCR from SAM and flower primordia up to stage 5 revealed that *LFY* RNA levels were downregulated by approximately 60% in *shr-2* compared to WT (Suppl Fig. 12D). We further tested the LFY protein level and expression pattern using *pLFY:GFP-LFY* reporter (previously named *pLFY::GLFY*) (Wu *et al*., 2003). In WT, *LFY* was not expressed in the SAM, but in flower primordia from stage 3 onwards. At later stages, *LFY* remained expressed in the adaxial side of sepals, and in the 2nd and 3rd floral whorls. In *shr-2* mutants, LFY-GFP fluorescence was barely visible at early stages and strongly decreased at later stages, compared to WT (Suppl Fig. 12B-C). This indicates that *SHR* function is required for normal *LFY* expression patterns. However, analysis of *SHR* and *SCR* expression in *lfy-12* mutants revealed no obvious differences compared to WT (Suppl Fig. 13A-D), and we therefore conclude that *LFY* does not feed-back to the *SHR-SCR* module. In the *lfy-12* null-mutant, leaf-like bracts are generated from the inflorescence meristem and formation of flower meristems is delayed. Most floral organs are leaf-like and not arranged in regular whorls. Double mutants of *lfy-12; shr-2* were strongly retarded in growth and displayed dramatically enhanced floral primordium initiation defects (Suppl Fig. 12E-L). We conclude from this analysis that the *SHR-SCR* module promotes *LFY* expression, and that it also promotes floral meristem development in a *LFY*-independent manner.

### *SCL23* genetically interacts with the *SHR-SCR* pathway and contributes to SAM size maintenance

The closest homologue of SCR is the GRAS-family TF SCARECROW-LIKE23 (SCL23). We investigated the expression pattern of *SCL23* in the SAM using the transcriptional reporter *pSCL23:H2B-YFP*, and the functional translational reporter line *pSCL23:SCL23-YFP (Long et al., 2015a)*. *pSCL23:H2B-YFP* was expressed in the L3 of the SAM and floral meristems, and in floral organ primordia (Fig. 11A and C). The translational reporter *pSCL23:SCL23-YFP* showed a wider expression pattern than the transcriptional reporter (Fig. 11B and D), which was suggestive of SCL23 protein mobility. To test if the wider expression pattern of the translational reporter line is due to the presence of additional transcriptional control regions within the coding sequence of *SCL23*, we expressed SCL23-mVenus under the *CLV3* promoter, which confines the expression exclusively to the CZ. We observed SCL23 spreading from the expression domain in the CZ into the surrounding cells, consistent with high SCL23 protein mobility in the SAM (Fig. 11E and F). We then analysed co-expression of SCL23 and SCR to investigate the regulatory dynamics among *SCL23*, *SCR* and *SHR*. A dual reporter line with *pSCR:SCR-RFP* and *pSCL23:SCL23-YFP* revealed that the expression patterns of both genes are in most cases mutually exclusive, with SCL23 in a central domain of the SAM and in the inner cell layers of primordia, while SCR is confined to the PZ, the L1 of the CZ and the outer cell layers of lateral organ primordia (Fig. 11G, I and K). Some cells expressing both SCL23 and SCR were located deep inside lateral organ primordia (Fig. 11I and K (yellow arrowheads)). A similar coexpression analysis of *pSCL23:SCL23-YFP* with *pSHR:mScarlet-SHR* indicated an overlap of expression in the deep cell layers of lateral organ primordia (Fig. 11H, J and L (yellow arrowheads)). In *shr-2* mutants, *pSCL23:SCL23-YFP* remains only weakly expressed in the meristem centre (Fig. 11M and N), indicating that SHR is required to promote expression of *SCL23* in the SAM periphery. We then performed genetic interaction studies to understand the relationship between *SCL23*, *SCR* and *SHR* in the SAM. In *scl23-2* mutants, SAM size is significantly decreased compared to WT, or to the complemented line *pSCL23:SCL23-YFP/scl23-2* (Fig. 11O-R). Using the PlaCCI marker to analyze cell division patterns, we found that similar to *shr-2* and *scr-3* or *scr-4* mutants, the percentage of cells in G1 phase increased, indicating an extended G1 phase and delayed cell divisions in the SAM (Fig. 11S-U). Genetic analyses showed that *shr* mutants are epistatic to *scl23* mutants in double mutant combinations, and that *scr-3; scl23-2* double mutants and *shr-2; scr-3; scl23-2* triple mutants were additive with smaller plant rosettes and reduced plant stature (Suppl Fig. 14A-H’) (Yoon *et al*., 2016). To further analyse *SCL23* function, we generated transgenic Arabidopsis plants overexpressing *SCL23* from the *pUBIQ10* (*pUBIQ10:SCL23-mVenus*) or *pRPS5A* (*pRPS5A*:*SCL23-mVenus*) promoter. Both *UBIQ10* and *RPS5A* promoters drive high-level and widespread expression of the fused reporter gene. However, we observed SCL23-mVenus fluorescence confined to the lateral organ primordia and not in the meristem center, where endogenous SCL23 is normally expressed (Suppl Fig. 15C-D). We found a similar scenario when we tried to overexpress *SHR* or *SCR* in the SAM (Suppl Fig. 15A-B), implying tight regulatory controls acting in the different domains in the SAM preventing excessive protein accumulation. However, excessive accumulation of these transcription factors was observed outside their normal expression domains (Suppl Fig. 15A-E). We propose that SCL23, SHR and SCR are under translational or post-translational control, for example due to the localised presence of miRNAs (Llave *et al*., 2002), and that meristem cells can therefore express only a set maximum amount of these GRAS proteins. We also conclude that SHR controls expression of both SCL23 and SCR in the periphery of the SAM.

**Fig. 11:**
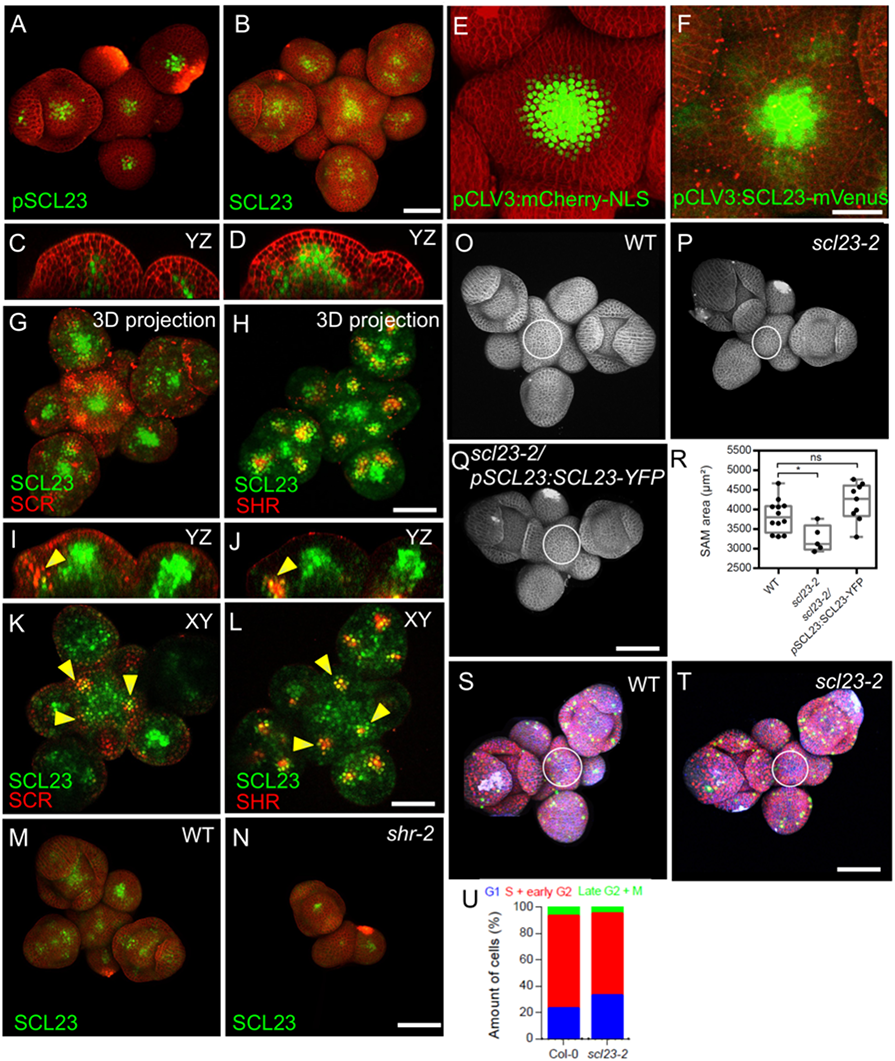
SCL23, SCR and SHR proteins show spatially different patterns but perform similar functions in the SAM. **(A and B)** Representative 3D projection of SAMs at 5 weeks after germination expressing *pSCL23:H2B-YFP* reporter (green) (n=5) **(A)** and *pSCL23:SCL23-YFP* reporter (green) (n=25) **(B)**. Cell walls were stained with PI (red). Scale bar represents 50 µm. **(C and D)** Longitudinal optical sections of **(A)** and **(B)** respectively. **(E and F)** Representative 3D projection of SAMs at 5 weeks after germination expressing *pCLV3:mCherry-NLS* reporter (green) (n=4) **(E)** and the *pCLV3:SCL23-mVenus* reporter (green) (n=13) **(F)**. Cell walls were stained with PI (red). Scale bar represents 50 µm. **(G and H)** Representative 3D projection of SAMs at 5 weeks after germination meristems coexpressing *pSCL23:SCL23-YFP* reporter (green) and *pSCR:SCR-RFP* reporter (red) (n=6) **(G)** and *pSCL23:SCL23-YFP* reporter (green) and *pSHR:mScarlet-RFP* reporter (red) (n=7) **(H)**. Scale bar represents 50 µm. **(I and J)** Longitudinal optical sections of **(G)** and **(H)** respectively. **(K and L)** Transversal optical sections of **(G)** and **(H)** respectively. Yellow arrowheads in **(I)** and **(K)** indicates the region where both *pSCL23:SCL23-YFP* reporter (green) and *pSCR:SCR-RFP* reporter (red) expression overlap. Yellow arrowheads in **(J)** and **(L)** indicates the region where both *pSCL23:SCL23-YFP* reporter (green) and *pSHR:mScarlet-SHR* reporter (red) expression overlap. **(M and N)** Representative 3D projection of SAMs at 5 weeks after germination expressing *pSCL23:SCL23-YFP* reporter (green) in WT (n=14) **(M)** and *shr-2* mutant (n=4) **(N)**. Cell walls were stained with PI (red). Scale bar represents 50 µm. **(O-Q)** Representative 3D projection of SAMs at 5 weeks after germination from WT (n=12) **(O)**, *scl23-2* mutant (n=5) **(P)** and *scl23-2/pSCL23:SCL23-YFP* (n=9) **(Q)**. **(R)** Quantification of SAM size at 5 weeks after germination from Col-0 (n=12), *scl23-2* mutant (n=5) and *scl23-2/pSCL23:SCL23-YFP* (n=9). Cell walls were stained with PI (gray). Scale bar represents 50 µm. **(S and T)** Representative 3D projection of SAMs at 5 weeks after germination from WT (n=11) **(S)** and *scl23-2* (n=5) **(T)** coexpressing the three PlaCCI markers. *pCDT1a:CDT1a-eCFP* reporter (blue), *pHTR13:pHTR13-mCherry* reporter (red) and *pCYCB1;1:NCYCB1;1-YFP* reporter (green). Cell walls were stained with DAPI (gray). Scale bar represents 50 µm. **(U)** Quantification of different cell cycle phases of SAM expressing PlaCCI in WT (n=6) and *scl23-2* (n=5). Asterisks indicate a significant difference (^∗^p < 0.01: Statistically significant differences were determined by Student’s *t*-test, ns= no significant difference).

### SCL23 acts together with WUS to maintain stem cell homeostasis in the SAM

The *GRAS* family transcription factors HAM1 and HAM2 were shown to physically interact with WUS to promote shoot stem cell proliferation and repress *CLV3* expression in the OC (Zhou *et al*., 2015). Since *SCL23* and *WUS* expression strongly overlap in the OC and rib meristem (Fig. 12A-B and D-E), we asked if SCL23, like HAM1/2, could also interact with WUS. We tested via FRET for *in vivo* interactions of WUS-FP and SCL23-FP fusions after inducible transient expression in *Nicotiana benthamiana* leaves. SCL23-mVenus interacted with WUS-mCherry with a FRET efficiency similar to the FRET-positive control, a nuclear localized mVenus-mCherry fusion protein (Fig. 12C). SCL23 showed only a weak tendency to form homomers, assayed by measuring FRET between SCL23-mVenus and SCL23-mCherry (Fig. 12C). SCL23-mVenus with the unrelated transcription factor AS2-mCherry served as negative control (Fig. 12C). Given its interaction with WUS and its developmental role in the SAM, we asked if SCL23 expression is also subject to regulation by the *CLV* signaling pathway. We first observed the expression pattern of *SCL23* and *CLV3*. A dual reporter line with *pSCL23:SCL23-YFP* and *pCLV3:NLS-mCherry* showed that *SCL23* and *CLV3* were expressed in largely complementary patterns in the SAM (Fig. 12A-B and D-E). SCL23 was expressed in the rib meristem, but absent from L1 and L2 layers in the meristem center, while *CLV3* is highly expressed in the L1 and L2 layers and to a lesser level in the L3 layer (Fig. 12D-E). When introduced into the *clv3-9* mutant background, *pSCL23:SCL23-YFP* expression was strongly expanded and extended throughout the enlarged and sometimes fasciated meristems, similar to the altered expression domain of *WUS* in strong *clv*-mutants (Fig. 12F-I). This supports the hypothesis that *SCL23* expression is, like *WUS*, negatively regulated by the *CLV* signalling pathway. To study the developmental role of *SCL23* and *WUS* interaction, we conducted genetic interaction studies. While the SAM of *scl23-2* single mutant was smaller than WT (Fig. 11R), the null allele *wus-am* had terminated SAM development at an early seedling stage, but continued to initiate new shoot meristems from axillary positions (Suppl Fig. 16C). The *wus-am; scl23-2* double mutants showed an enhanced phenotype with earlier arrest of SAM development (Suppl Fig. 16D). Hypomorphic *wus-7* mutants form inflorescence meristems similar to WT (Suppl Fig. 16E,E’,E’’,F,F’,F’’,G,G’ and G’’) (Zhou et al., 2015), while *wus-7; scl23-2* double mutants display premature termination of floral meristems (Fig. 15I-I’-I’’). These genetic data support our hypothesis that SCL23 and WUS function together in shoot meristem stem cell maintenance. Taken together, the colocalization and interaction between SCL23 and WUS along with the genetic data strongly suggest that SCL23 and WUS function as partners in SAM maintenance.

**Fig. 12:**
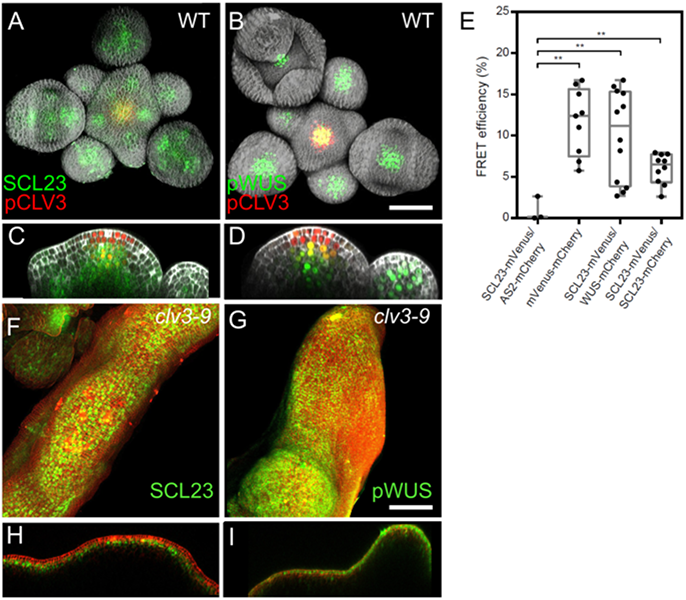
SCL23 and WUS physically interact and are negatively regulated by CLV signalling (A and B) Representative 3D projection of SAMs at 5 weeks after germination coexpressing *pSCL23:SCL23-YFP* (green) and *pCLV3:mCherry-NLS* reporter (red) (n=3) **(A)** and *pWUS:3xVenus-NLS* reporter (green) and *pCLV3:mCherry-NLS* reporter (red) (n=6) **(B)**. Cell walls were stained with DAPI (gray). Scale bar represents 50 µm. **(C and D)** Longitudinal optical sections of **(A)** and **(B)** respectively. **(E)** FRET efficiency measured in epidermis cells of *N. benthamiana* between SCL23-mVenus and WUS-mCherry (n=12) or SCL23-mCherry (n=10), compared with the negative control SCL23-mVenus and AS2-mCherry (n=3) and positive control mVenus-mCherry (n=9). **(F and G)** Representative 3D projection of shoot apical meristems at 5 weeks after germination expressing *pSCL23:SCL23-YFP* reporter (green) in *clv3-9* mutant (n=3) **(F)** and *pWUS:3xVenus-NLS* reporter (green) in *clv3-9* mutant (n=3) **(G)**. Cell walls were stained with PI (red). Scale bar represents 50 µm. **(H and I)** Longitudinal optical sections of **(F)** and **(G)** respectively. Asterisks indicate a significant difference (^∗∗^p < 0.001: Statistically significant differences were determined by Student’s *t*-test).

## Discussion

To better understand the mechanisms that form and maintain the multiple functional domains within the SAM, we asked if the well-characterized SHR-SCR-SCL23-JKD gene regulatory network responsible for ground tissue patterning in the RAM plays a similar role in the SAM. (Scheres et al., 1995, Long et al., 2015a, Long et al., 2015b). Using quantitative live-cell imaging together with molecular and genetic approaches, we found that all four TFs are expressed in the SAM and together their expression covers all functional domains of the SAM in a combination of complementary and overlapping domains. Auxin-triggered primordia initiation in the peripheral zone of the SAM activates the SHR network, which promotes cell cycle progression and, at the same time, communicates with the stem cell regulatory system at the center of the meristem through direct interaction with the WUS-CLV3 feedback loop.

### SHR, SCR, SCL23 and JKD control SAM size and organ primordia initiation

Plants mutated in the three GRAS genes (*SHR*, *SCR* or *SCL23*) had smaller SAMs than the WT caused by a decrease in cell proliferation (Fig. 1E-H, I-L and L-P; Fig. 11O-U). *shr* and *scr* mutants also displayed a delay in primordia formation suggesting that the *SHR, SCR* and *SCL23* genes promote proliferation of stem cells within the central zone and primordia initiation at the peripheral zone (Fig. 1M; Suppl Fig. 1A’-D’). These findings are consistent with the growth-promoting activities of *SHR* and *SCR* in leaves and roots (Dhondt *et al*., 2010, Long et al., 2015a, Scheres et al., 1995). In contrast, *jkd* mutants showed an increase in the SAM size indicating that JKD might restrict growth in the shoot by interfering with the SHR-SCR complexes, similar to what had been previously shown in roots (Fig. 6A-D) (Long et al., 2015b, Welch et al., 2007).

### Transcriptional profiles and interdependencies of SCR, SCL23 and JKD in the SAM

Analysis of expression patterns showed that SHR, SCR, SCL23 and JKD are distributed across all functional domains of the SAM, and mutant analyses suggest that the SHR network is required for SAM function and organogenesis. We found both SHR and SCL23 to be mobile in the SAM with SCL23 exhibiting hypermobility, moving from the center to the periphery of the SAM, similar to what was previously shown in the RAM (Fig. 3C-D; Fig. 11A-F). SCL23 could thereby mediate communication between the CZ and PZ, and coordinate stem cell proliferation in the center with lateral organ formation at the periphery via the conserved SHR signalling pathway. SHR acts as master regulator in the network, since all combinations between *shr, scr* and *scl23* mutants displayed the *shr* phenotype (Suppl Fig. 14A-H’) (Yoon et al., 2016). Previous studies had shown that SCR and SCL23 restrict the mobility of SHR, probably by complex formation, and both SCR and SCL23 are transcriptionally regulated by SHR in the RAM. Similarly, we found here that the expression of *SCR* and *SCL23* was down-regulated in the *shr* mutant organ primordia (Fig. 5D-E and H-I; Fig. 11M-N) and SHR is more restricted to the nucleus in the presence of SCR compared to its absence (Fig. 5A, 5A inset and F). This could be explained by the nuclear retention of SHR through its interaction with SCR to restrict SHR movement and activate SCR expression. Together, these results suggest that SHR regulates the expression of SCR and SCL23, which in turn restrict SHR mobility. SCL23 and SCR have been previously shown to function in a mutually antagonistic manner in the RAM and act redundantly to specify endodermal cell fate (Kim et al., 2017, Long et al., 2015a). Our data also showed that the expression patterns of SCL23 and SCR are complementary in the center of the SAM, but overlap in the primordia where SHR and JKD are also expressed (Fig. 6I; Fig. 11G-K).

### SHR, SCR, SCL23 and JKD interact to form protein complexes in the SAM

In root CEI, a SHR-SCR-JKD complex controls the expression of CYCD6;1 (Suppl Fig. 17A-G arrow heads) and triggers an asymmetric cell division that leads to the formation of endodermis and cortex (Cui et al., 2007, Sozzani *et al*., 2010, Long et al., 2015b, Long et al., 2017). Using *in vivo* FRET-FLIM, we now showed that SHR, SCR and JKD physically interact in the SAM (Fig. 4E-F; Fig. 6H). Furthermore, expression analysis of targets of the SHR-SCR-JKD complex demonstrates that CYCD6;1 expression perfectly overlaps with that of SHR, SCR and JKD in the inner cells of the lateral primordia (Fig. 7J-L). *shr* mutants showed a down-regulation of *CYCD6;1* and a decrease in the number of cell layers at the PZ (Fig. 7A-I). We conclude that the SHR-SCR-SCL23-JKD complex acts as a positive regulator of cell proliferation within the PZ leading to lateral organ initiation and outgrowth.

### SHR regulates the initiation of flower primordia in an auxin dependent manner

The expression patterns of SHR closely resemble that of PIN1 and MP during organ primordia development, and also in the RAM (Fig. 3C; Suppl Fig. 6A-C and E-F). We now show that SHR expression responds to changes in auxin distribution in the SAM and that the addition of exogenous synthetic auxin enhances its expression (Fig. 8A-D and H-K). Moreover, we found that MP-mediated auxin signalling positively regulates *SHR* expression (Fig. 9A-F and K-P). SHR has been previously proposed to be a putative downstream target of MP, as SHR expression was found to be down regulated in *mp* mutant embryo (Möller et al., 2017). Our genetic and expression analysis now revealed that MP is a direct regulator of SHR expression. The two AuxRE motifs in the *SHR* promoter were found to be essential for the response to MP in our transient expression experiments (Suppl Fig. 8A-J; Suppl Fig. 9A-J; Suppl Fig.10A-H). These results suggest that the AuxRE motifs enable auxin-mediated regulation of *SHR* expression and both motifs have roughly equal contribution in this transcriptional regulation. Thus, the SHR network regulates organogenesis in the SAM downstream of MP in response to auxin accumulation at the meristem periphery, and thereby plays a crucial role for lateral organ formation. Moreover, the altered expression of both *pDR5v2:3xYFP-N7* and R2D2 auxin reporters in *shr* and *scr* mutants (Fig. 2A-G) with fewer auxin maxima indicates that *SHR* and *SCR* contribute to a positive feed-back loop with auxin.

### A WUS-SCL23 heteromeric complex maintains the stem cell niche in the SAM

Unlike the other genes in the SHR network, *SCL23* was expressed in a broad pattern in the OC, PZ and in lateral organ primordia (Fig. 11A-D). SCL23 overlapped with WUS expression in the OC, but is excluded from the CZ (Fig. 12D-E). WUS protein migrates from the OC to the CZ where stem cells are located and activates the expression of its negative regulator CLV3 (Daum et al., 2014, Yadav et al., 2011). The resulting WUS-CLV3 negative feedback loop maintains the balance between stem cell numbers at the CZ and differentiation rates of stem cell descendents at the PZ (Daum et al., 2014). Null mutations in *WUS* cause premature termination of the SAM (Suppl Fig. 16C) (Laux et al., 1996). Two GRAS TFs, HAM1 and HAM2, are expressed in the OC, physically interact with WUS and inhibit WUS from activating *CLV3* expression in the OC. At the meristem tip and in the CZ, *HAM1* and *HAM2* transcripts are downregulated due to miRNA171, which is expressed from the L1 and diffuses downward into deeper meristem layers. In consequence, WUS lacks its binding partners HAM1 and HAM2 only in the top layers of the meristem where miRNA171 levels are high, and WUS can then promote stem cell fate and activate CLV3 expression (Zhou et al., 2015, Zhou et al., 2018). SCL23 physically interacts with WUS, and SAMs lacking SCL23 activity are smaller, indicating that the SCL23-WUS complexes promote stem cell activity (Fig. 11R and Fig. 12C;). However, SCL23 appears to lack the recognition sequences for regulation by miRNA171, but is itself transcriptionally regulated by CLV3 signalling. Thus, two types of GRAS transcription factors modulate WUS activity: (i) the HAMs, due to their regulation by miRNA171, position the domain of WUS activity relative to the SAM surface (L1); (ii) SCL23 sets the WUS domain in relation to the CLV3 domain and, since SCL23 is regulated by the auxin-controlled SHR network, in relation to lateral organ primordia formation in the PZ.

### A SHR-SCR-SCL23-JKD regulatory network coordinates SAM activities

We propose that a gene regulatory network consisting of mobile transcription factors, together with auxin, controls SAM functions (Fig. 13A-B). *SCL23* gene expression is confined by CLV3 signalling to the OC, where SCL23 forms a complex with WUS to maintain stem cell homeostasis. SCL23 might work in a similar way as its closest homologues, the HAM proteins, which act as WUS partners to control shoot stem cell proliferation. The hypermobile SCL23 protein acts as a signal from the OC to the PZ (Fig. 13B). In the PZ, auxin promotes SHR expression via MP. SHR then physically interacts with SCR, SCL23 and JKD to activate the expression of *CYCD6;1* and promote cell division, leading to lateral organ primordia outgrowth.

**Fig. 13:**
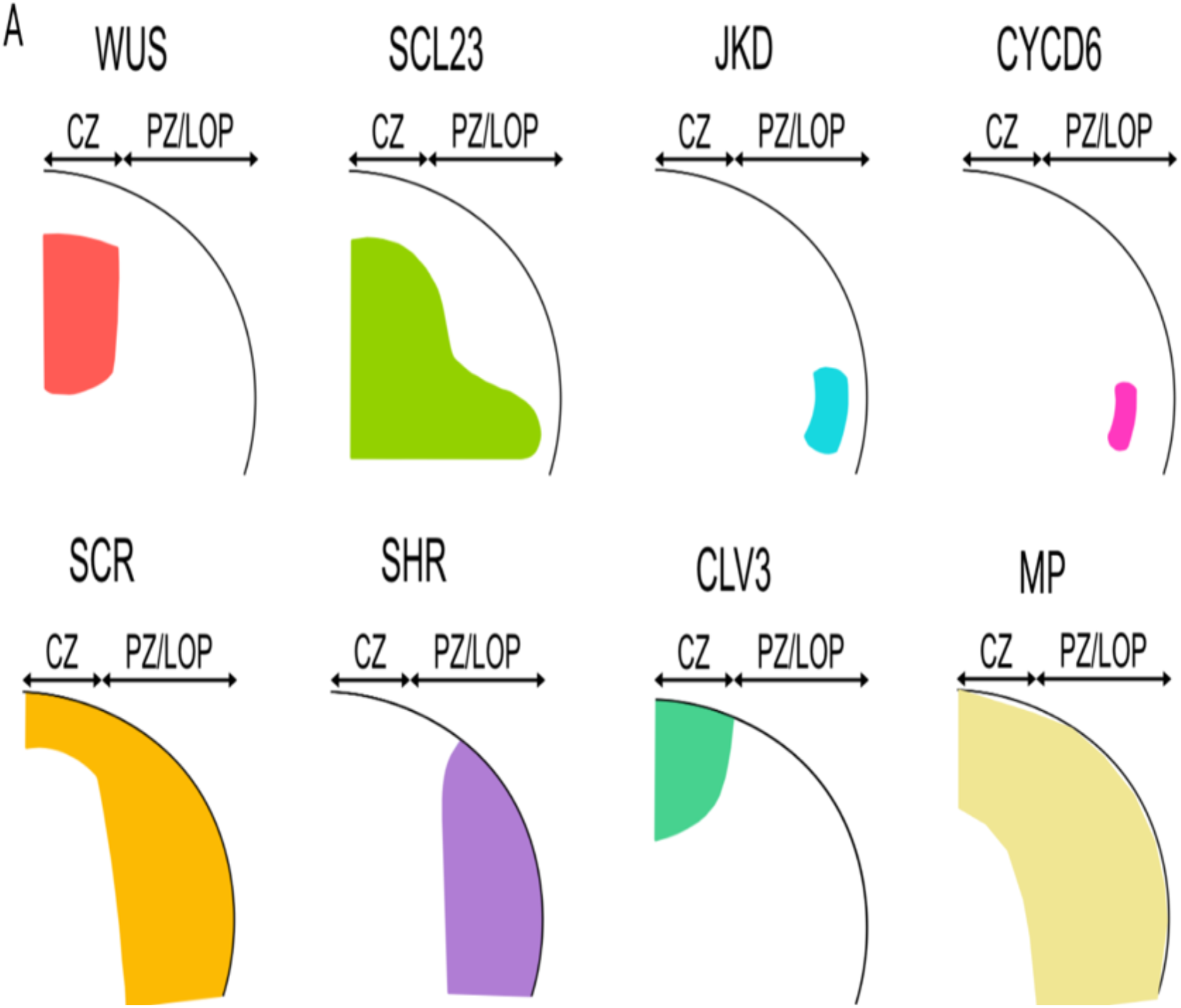
Proposed model for SHR-SCR-SCL23-JKD regulatory network function in the SAM. **(A)** Schematic representation of observed expression patterns of *WUS*, *SCL23, JKD, SCR, SHR, CLV3*, *MP* and *LFY* showing typical expression domains in the SAM. **(B)** Schematic molecular models for SHR-SCR-SCL23-JKD gene regulatory network function in the SAM, genes with genetic and/or biochemical interactions are indicated. Lines with arrows depict positive regulation, and with bars depict negative regulation. Dashed lines with arrows depict intercellular protein movement. Overlap between circles describe protein–protein interactions. In the organizing center (OC), SCL23 interacts with WUS to maintain stem cell homeostasis in the central zone (CZ). Like WUS, SCL23 is subject to regulation by CLV3 signalling. SCL23 is hypermobile and moves to the peripheral zone (PZ), where *SHR* transcription is promoted by auxin through MP. In the PZ, SHR activates *SCR* transcription. The SHR-SCR-SCL23-JKD protein complex promotes cell cycle progeression and induces periclinal cell divisions that lead to the outgrowth of lateral organ primordia through the activation of *CYCD6;1*. SHR also regulates the expression of *LFY* within the corpus in the PZ/LOP leading to lateral organ initiation. CZ=Central zone, OC=Organizing center, PZ/LOP=Peripheral zone/lateral organ primordia.

## Acknowledgements

We thank Jan Lohmann, Dolf Weijers, Francois Parcy and Doris Wagner for providing seeds. We are also grateful to the CAi team at HHU, especially Sebastian Hänsch for support with imaging methods, and Patrick Blümke, Jan Maika, Jenia Schlegel, Maike Breiden, Meik Thiele, Karine Gustavo Pinto, Svenja Augustin, Carin Theres, Silke Winters, Vivien Strotmann, Rebecca Drisch, Madhumita Narasimhan, Vicky Howe, Pavithran Narayanan and Cornelia Gieseler for diverse contributions.

Research in the lab of R.Simon is supported by the DFG through the Cluster of Excellence on Plant Sciences (CEPLAS, EXC2048), CRC1208 and RTG CSCS.

Research in the lab of Crisanto Gutierrez is supported by the Grant ERC-2018-AdG_833617 (European Union, H2020) and Grant RTI2018-094793-B-I00 (from Spanish Ministry of Science and Innovation, and FEDER).

## Materials and methods

### Plant material and growth conditions

*Arabidopsis thaliana* plants were grown on soil in climate chambers under long day (LD) conditions (16 h light / 8 h dark) at 21 °C. Most plants used in this study were in the Columbia (Col-0) background, except for: *scr-4* (WS) (Fukaki *et al*., 1998) and *wus-*7 (Graf *et al*., 2010) in the *Landsberg erecta* (L.er.) background (Tab. 8 and 9).

### Plasmid construction and plant transformation

The entry plasmids in this study (Tab. 5) were generated by gDNA or cDNA sequences that were amplified with Phusion High-Fidelity PCR polymerase using primers described in Tab. 1. All destination vectors in this study (Tab. 6 and Tab. 7) were generated using the GreenGate cloning system (Lampropoulos *et al*., 2013). Entry plasmid containing SHR promoter sequence (2.5 KB) was used as a template to create different promoter mutants using the QuikChange II kit according to manufacturer’s protocol (Agilent Technologies). Mutagenic mismatch primers are listed in Tab. 2. all the clones were confirmed by sequencing. *Arabidopsis thaliana* wildtype (Col-0) plants were transformed with *Agrobacterium tumefaciens* (strain GV3101 pMP90 pSoup) containing the respective destination vectors using the floral dipping method (Clough & Bent, 1998). Transgenic plants were initially selected on the appropriate antibiotic in the T1 generation. Homozygous T2 plants were identified through confirmation in the T3 generation. Several independent lines were analysed and a representative line was selected for further work. *Nicotiana benthamiana* plants were grown 4 – 5 weeks in the greenhouse and subsequently used for transient leaf epidermis cell transformation.

**Tab. 1:** Enzymes used in this study

| Enzyme | Producer |
| --- | --- |
| BSA1 (ECORI) | Thermo Fisher Scientific, Braunschweig, Germany |
| PfuUltra High-Fidelity DNA polymerase | Agilent, Santa Clara, USA |
| Phusion® High-Fidelity DNA Polymerase | Thermo Fisher Scientific, Braunschweig, Germany |
| T4-Ligase | Thermo Fisher Scientific, Braunschweig, Germany |
| Taq-DNA-Polymerase | Made in the lab according to (Pluthero, 1993) |
| Universal SYBR® Green Supermix | Bio-Rad |

**Tab. 2:**
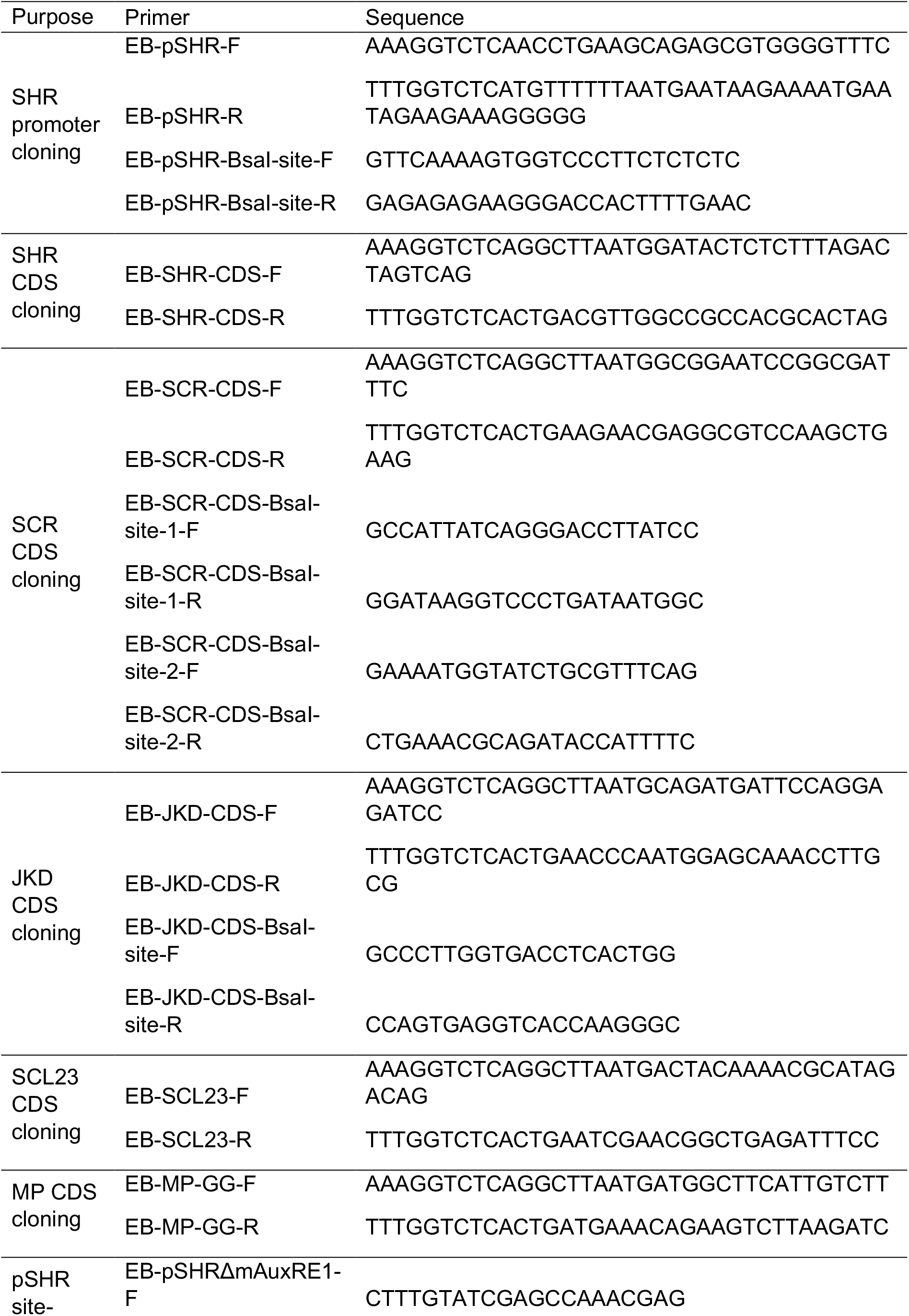
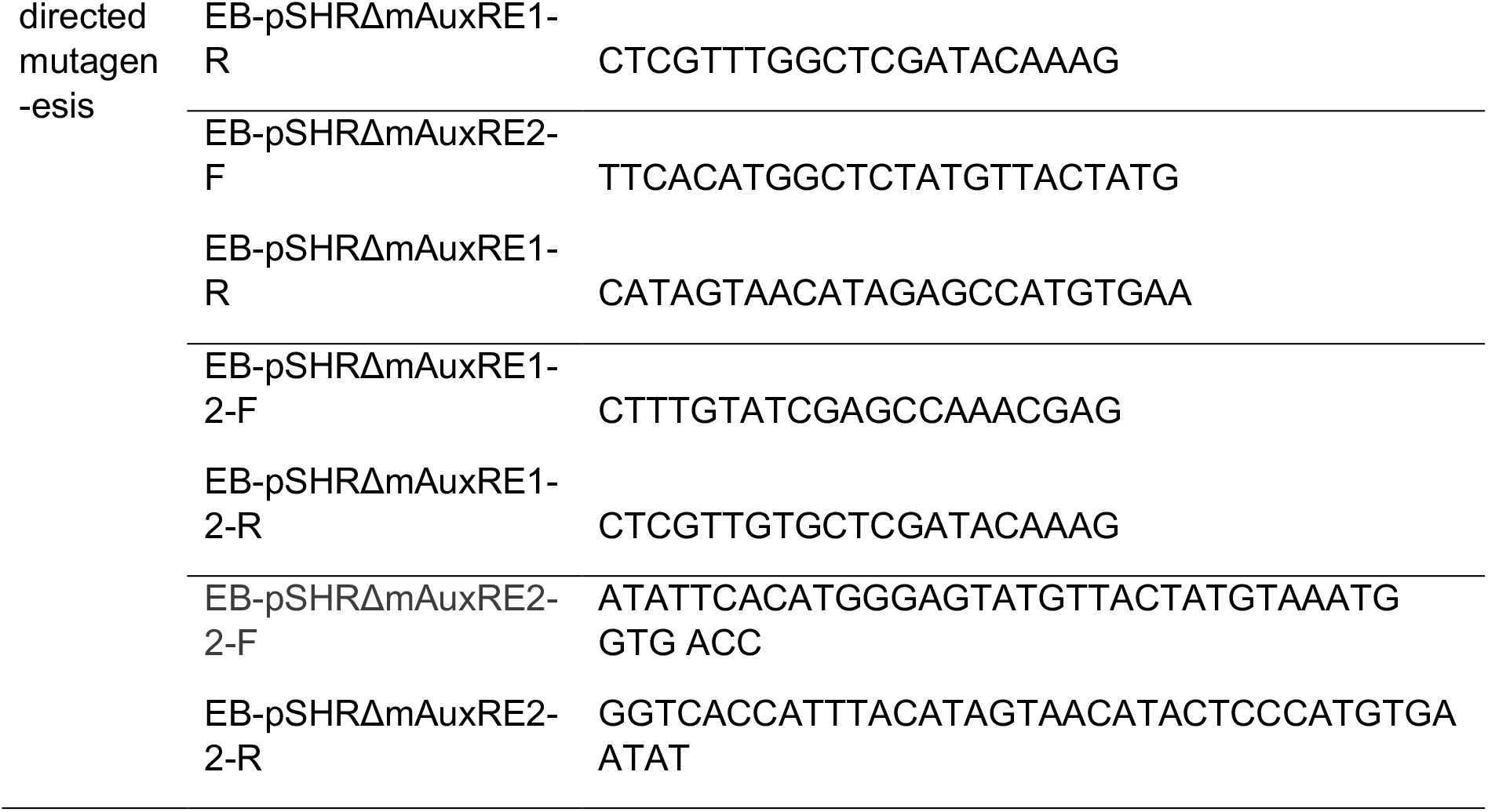
Primers used for cloning

**Tab. 3:**
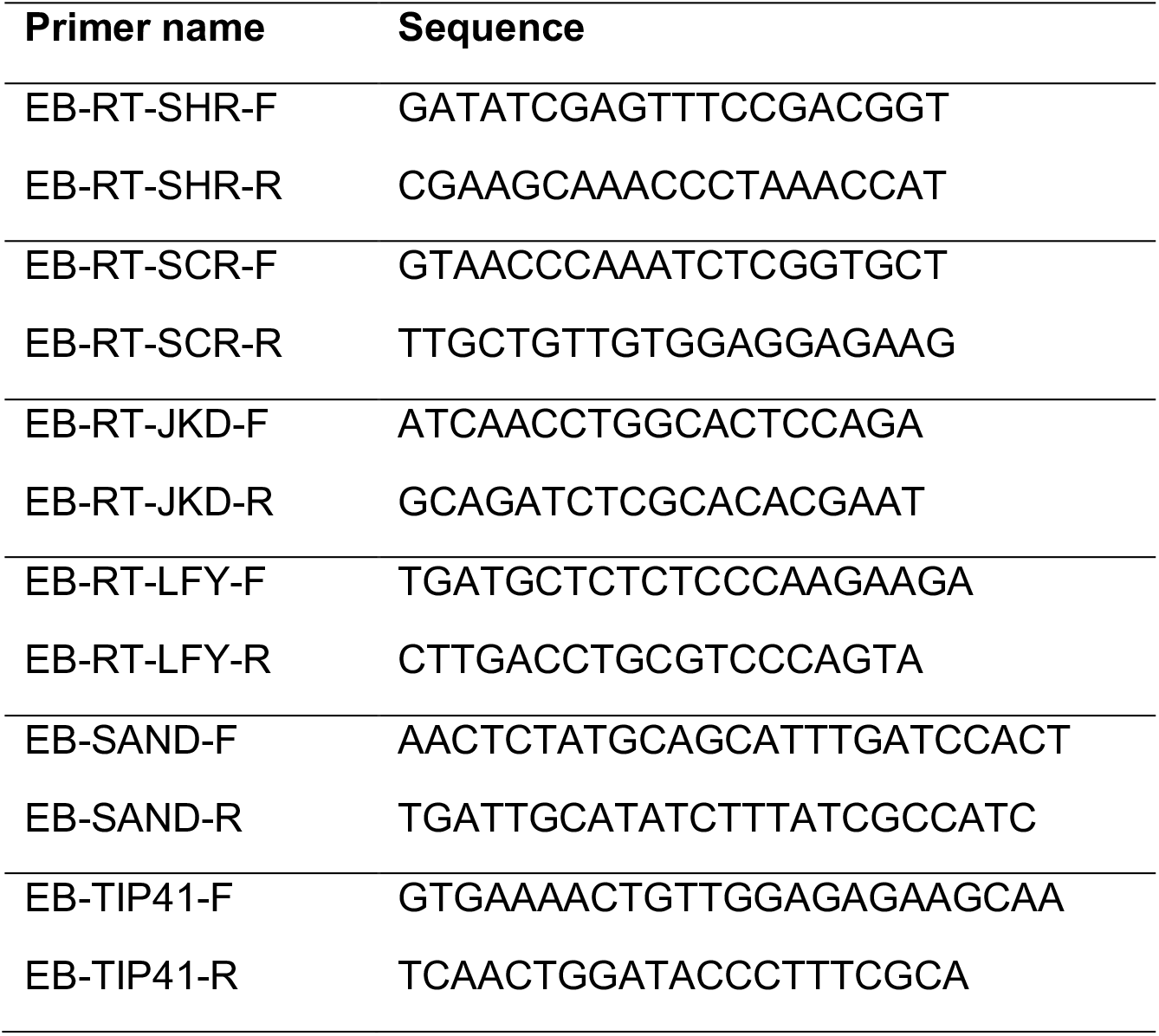
Primers used for qRT-PCR

**Tab. 4:**
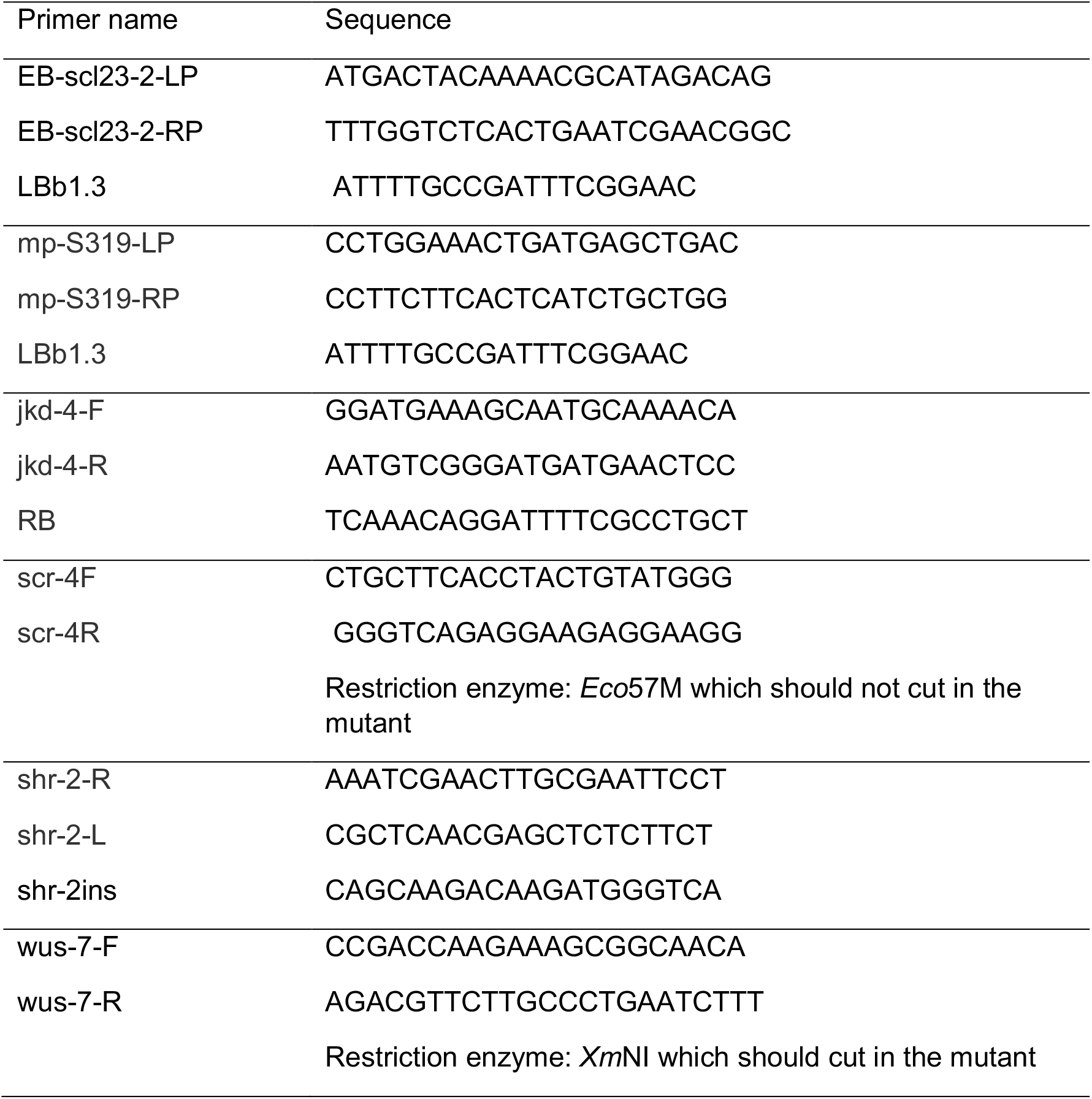
Primers used for genotyping

**Tab. 5:** Entry plasmids used in this study

| Name | Description | Backbone | Reference |
| --- | --- | --- | --- |
| pSHR | SHR promoter 2.5 Kb upstream from transcription start | pBlunt | This study |
| SHR CDS | SHR coding sequence | pGGC000 | This study |
| SCL23 CDS | SCL23 coding sequence | pGGC000 | This study |
| linker NLS (pGGD007) | NUCLEAR LOCALIZATION SIGNAL | pGGD000 | Lampropoulos et al., 2013 |
| mVenus (pRD43) | mVenus | pGGD000 | Rebecca Burkhart |
| FLUC | Firefly luciferase | pGGC000 | Greg Denay |
| mVenus (pRD42) | mVenus | pGGC000 | Rebecca Burkhart |
| GUS (pGGC051) | E. coli β-GLUCURONIDASE | pGGC000 | Lampropoulos et al., 2013 |
| MP CDS | MP coding sequence | pGGC000 | This study |
| 35S promoter (pGGA004) | Cauliflower mosaic virus 35S promoter | pGGA000 | Lampropoulos et al., 2013 |
| RPS5A promoter (pGGA012) | RIBOSOMAL PROTEIN 5A promoter | pGGA000 | Lampropoulos et al., 2013 |
| UBIQ10 promoter (pGGA006) | UBIQUITIN10 promoter | pGGA000 | Lampropoulos et al., 2013 |
| d-dummy (pGGD002) | d-dummy | pGGD000 | Lampropoulos et al., 2013 |
| tCLV3 | CLV3 terminator 1257 bp downstream of transcription stop | pGGE000 | Jenia Schlegel |
| UBQ10 terminator (pGGE009) | UBQ10 terminator | pGGE000 | Lampropoulos et |
| BastaR (pGGF008) | pNOS:BastaR:tNOS | pGGF000 | Lampropoulos et al., 2013 |
| GR (pRD64) | Hormone-binding domain of the glucocorticoid receptor | pGGD000 | Rebecca Burkhart |
| pCLV3 | CLV3 promoter 1480 bp upstream from transcription start | pGGA000 | Jenia Schlegel |
| omega-element (pGGB002) | Omega- element | pGGB000 | Lampropoulos et al., 2013 |

**Tab. 6:**
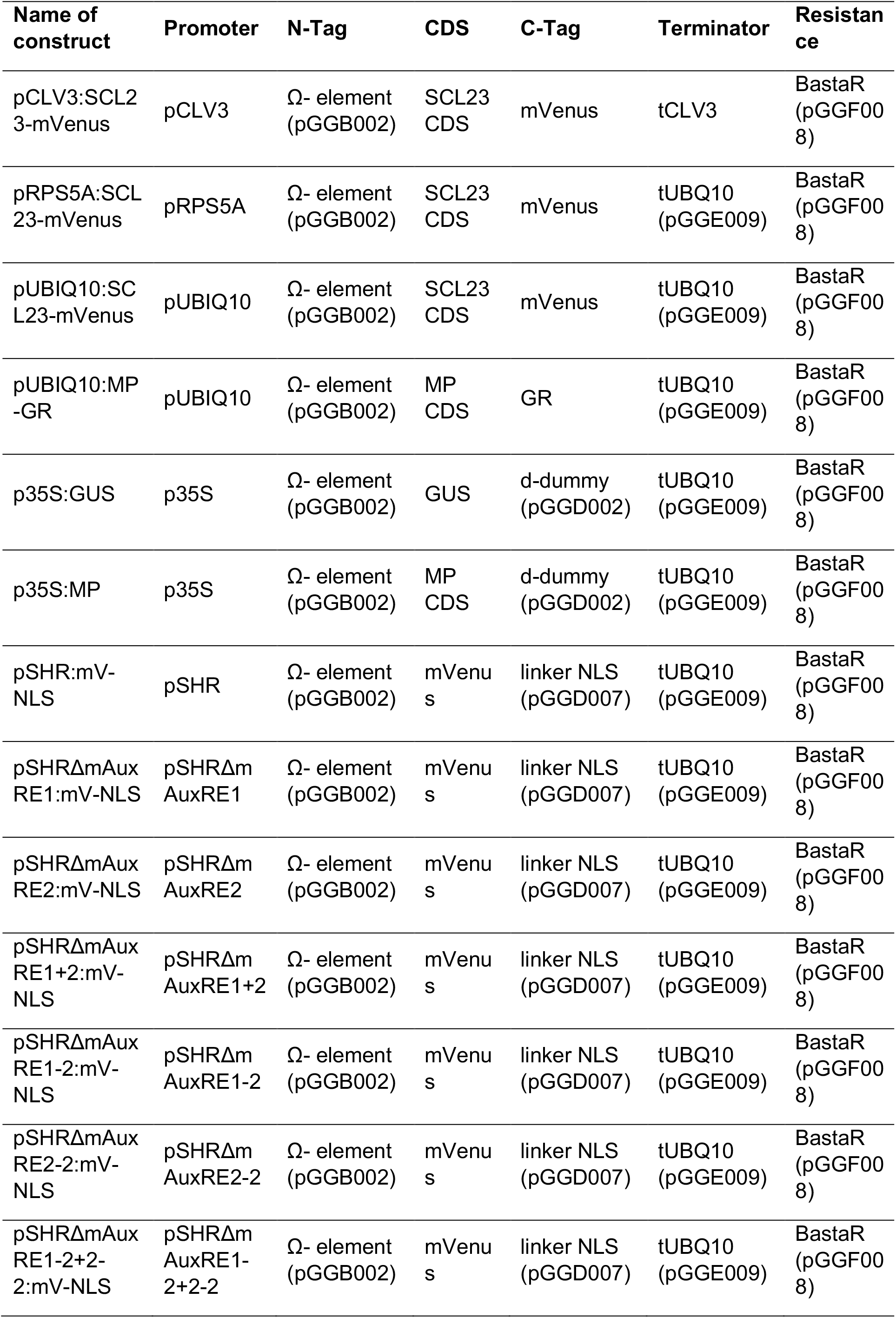
Destination plasmids used in this study

### Nicotiana benthamiana infiltration

*Agrobacterium tumefaciens* strains (strain GV3101 pMP90 pSoup) harbouring relevant reporter constructs were cultured overnight with shaking at 28 °C in 5 ml dYT (double Yeast Tryptone, 1.6 % w/v tryptone, 1 % w/v yeast extract, 0.5 % w/v NaCl) with appropriate antibiotics. Cell cultures were adjusted to an optical density (OD_600_) of 0.3 and then harvested by centrifugation at 4000xg for 10 min. The pellet was resuspended in infiltration buffer (5% w/v sucrose, 150μM acetosyringone, 0.01% v/v Silwet) and incubated for 2h at 4°C. For coexpression of two transgenes, the corresponding transformed *A. tumefaciens* were mixed equally. The resuspensions were infiltrated into the abaxial leaf surface of the 3-4 weeks old *N. benthamiana* using a needle-less syringe. Plants transformed with constructs under the control of the 35S promotor were used for analyses 4 days after infiltration.

### Chemical treatments

For hormone and dexamethasone treatments, plants were grown in soil. For RNA isolation experiments, *pSHR:SHR-YFP*, *pSCR:SCR-YFP* and *pCYCD6;1:GFP* expression analysis following auxin and auxin transport inhibitor treatment were performed by dipping 30-day-old plants inflorescence once in 10 µM (IAA or 2,4 D) once or with 100µM NPA twice (at 0h and 7 h). For Dexamethasone treatment *pUBIQ1:MP-GR* inflorescence meristems were treated with 10μM DEX only once and were imaged 5h after treatment.

### Promoter mVenus activity in *Nicotiana benthamiana*

*N. benthamiana* leaves were co-transformed with different *pSHR:mVenus-NLS* and the effector plasmids *p35S:GUS* or *p35S:MP* (Tab. 6). Four days after infiltration, the leaves were processed for further analysis using imaging with a Zeiss LSM780 using the same settings for all conditions.

### Luciferase assay in Nicotiana benthamiana

The different reporter constructs with firefly luciferase reporter (FLUC) under the control of the different versions of *SHR* promoter (Tab. 7) were co-infiltrated into *N. benthamiana* leaves together with the effector plasmids *p35S:GUS* or *p35S:MP* (Tab.6). Luciferase activities were measured four days after infiltration with the NightOwl luminescence system (Berthold). As substrate for the luciferase reaction, 5mM D-Luciferin potassium salt solution was used.

**Tab. 7:** Destination plasmids used for luciferase assay

| Name of construct | Promoter | N-Tag | CDS | C-Tag | Terminator | Resistance |
| --- | --- | --- | --- | --- | --- | --- |
| pSHR:FLUC | pSHR | Ω- element (pGGB002) | FLU C | d-dummy (pGGD002) | tUBQ10 (pGGE009) | BastaR (pGGF008) |
| pSHRΔmAuxRE1: FLUC | pSHRΔmA uxRE1 | Ω- element (pGGB002) | FLU C | d-dummy (pGGD002) | tUBQ10 (pGGE009) | BastaR (pGGF008) |
| pSHRΔmAuxRE2: FLUC | pSHRΔmA uxRE2 | Ω- element (pGGB002) | FLU C | d-dummy (pGGD002) | tUBQ10 (pGGE009) | BastaR (pGGF008) |
| pSHRΔmAuxRE1 +2:FLUC | pSHRΔmA uxRE1+2 | Ω- element (pGGB002) | FLU C | d-dummy (pGGD002) | tUBQ10 (pGGE009) | BastaR (pGGF008) |
| pSHRΔmAuxRE1- 2:FLUC | pSHRΔmA uxRE1-2 | Ω- element (pGGB002) | FLU C | d-dummy (pGGD002) | tUBQ10 (pGGE009) | BastaR (pGGF008) |
| pSHRΔmAuxRE2- 2:FLUC | pSHRΔmA uxRE2-2 | Ω- element (pGGB002) | FLU C | d-dummy (pGGD002) | tUBQ10 (pGGE009) | BastaR (pGGF008) |
| pSHRΔmAuxRE1- 2+2-2:FLUC | pSHRΔmA uxRE1- 2+2-2 | Ω- element (pGGB002) | FLU C | d-dummy (pGGD002) | tUBQ10 (pGGE009) | BastaR (pGGF008) |

**Tab. 8:** Mutants used in this study

| Lines | Reference |
| --- | --- |
| *scr-4* | Fukaki et al. 1998 |
| *jkd-4* | Welch et al., 2007 |
| *scr-3* | Fukaki et al., 1996b |
| *shr-2* | Nakajima et al. 2001 |
| *scl23-2* | Lee et al.,2008 |
| *wus-7* | Graf et al., 2010 |
| *wus-am* | From Jan lohmann |
| *mps-319* | Schlereth et al., 2010 |
| *mp-b4149* | Weijers et al., 2005 |
| *lfy-12* | Huala and Sussex,<br>1992 |
| *clv3-9* | Simon et al, 2003 |

**Tab. 9:** Transgenic lines used in this study

| **Lines** | **Plant Resistance** | **Reference** |
| --- | --- | --- |
| *pSHR:mV-NLS* | Basta | This study |
| *pSHRΔmAuxRE1:mV-NLS* | Basta | This study |
| *pSHRΔmAuxRE2:mV-NLS* | Basta | This study |
| *pSHRΔmAuxRE1+2:mV-NLS* | Basta | This study |
| *pSHRΔmAuxRE1-2:mV-NLS* | Basta | This study |
| *pSHRΔmAuxRE2-2:mV-NLS* | Basta | This study |
| *pSHRΔmAuxRE1-2+2-2:mV-NLS* | Basta | This study |
| *pCLV3:SCL23-mVenus* | Basta | This study |
| *pRPS5A:SCL23-mVenus* | Basta | This study |
| *pUBIQ10:SCL23-mVenus* | Basta | This study |
| *pUBIQ10:MP-GR* | Basta | This study |
| *PlaCCI* | Kanamycin | Desvoyes et al., 2020 |
| *pSHR:mScarlet-SHR* | - | This study |
| *pSHR:YFP-SHR* | Basta | Long et al, 2017 |
| *pSCR:SCR-RFP* | hyg | Long et al, 2017 |
| *pSCR:SCR-YFP* | basta | Long et al, 2017 |
| *pSHR:SHR-YFP* | Basta | Long et al, 2017 |
| *pJKD:mRFP-YFP* | norf | Long et al, 2017 |
| *pJKD:JKD-YFP* | Basta | Long et al, 2017 |
| *pSCL23:SCL23-YFP* | Kanamycin | Long et al. 2015a |
| *pSCL23:H2B-YFP* | Kanamycin | Long et al. 2015a |
| *pCYCD6;1:GFP* | Basta | Sozzani et al., 2010 |
| *pSCR:H2B-YFP* | Kanamycin | Heidstra et al, 2004 |
| *pPIN1:PIN1-GFP* | Kanamycin | Benkova et al., 2003 |
| *pSHR:nTdTOMATO* | - | Möller et al, 2017 |
| *pMP:MP-GFP* | Kanamycin | Schlereth et al, 2010 |
| *pLFY:LFY-GFP* | kanamycin | Wu et al., 2003 |
| *R2D2* | kanamycin | Liao et al, 2015 |
| *pDR5v2:3xYFP-N7* | Basta | Heisler et al, 2005 |
| *pWUS:3xVenus-NLS/pCLV3-mCherry-NLS* | Kanamycin | Pfeiffer et al, 2016 |
| *pCYCD1,1-GFP* | - | Forzani et al, 2014 |
| *pCYCD3,2-GFP* | - | This study |
| *pCYCD3,1-GFP* | - | This study |
| *pCYCD5,1-GFP* | - | This study |
| *pCYCD7,1-GFP* | - | This study |
| *pCYCD2,1-GFP* | - | This study |
| *pCYCD3,3-GFP* | - | Forzani et al, 2014 |
| *pCYCB1;1:CYCB1;1-GFP* | Kanamycin | Ubeda-Tomas et al., 2009 |

### Image acquisition and analysis

All confocal images were obtained by using a Zeiss LSM780 confocal microscope (40x water immersion objective, Zeiss C-PlanApo, NA1.2). For time series analysis, settings were established in the beginning on mock samples and were kept standard during the experiment. Shoot meristems were manually dissected by cutting of the stem, removing the flowers, and were stained with 1mg/ml DAPI or 5mM propidium iodide (PI). Individual populations of 2 to 20 plants were analyzed.

Green fluorescence was excited with an argon laser at 488 nm and emission was detected at 490 – 530nm, yellow was excited with an argon laser at 514 nm and emission was detected at 520 – 550nm, and red was excited with Diode-pumped solid state (DPSS) lasers at 514nm and detected at 570 – 650nm. PI was excited at 561nm by DPSS lasers and detected by PMTs at 590 – 650nm. DAPI was excited at 405nm with a laser diode and detected at 410 – 480nm.

### FRET-Acceptor-Photobleaching (APB)

*N. benthamiana* leaf epidermal cells were examined using a Zeiss LSM780 confocal microscope (40x Water immersion objective, Zeiss C-PlanApo, NA1.2). FRET was measured via mVenus fluorescence intensity increase after photobleaching of the acceptor mCherry (Bleckmann et al., 2010). The percentage change of the mVenus intensity directly before and after bleaching was analyzed as E_FRET_= (E_FRET_ = (mVenus_after_ – mVenus_before_)/mVenus_after_ x 100. All photobleaching experiments were performed in the nucleus. The displayed data were obtained from at least 3 independent experiments.

### FRET-FLIM interaction analysis in the SAM

In vivo FRET-FLIM experiments were measured using a Zeiss LSM780 confocal microscope (40x Water immersion objective, Zeiss C-PlanApo, NA1.2) equipped with a single-photon counting device (PicoQuant HydraHarp400) and a linear polarized diode laser (LDH-D-C-485). YFP donor fluorophores was excited with a 485nm (LDH-D-C-485, 32MHz) pulsed polarized diode laser. Excitation power was adjusted to 1,5μW. Emitted light was separated by a polarizing beam splitter and detected with a band-pass filter (520/35 AHF) by Tau-SPADs (PicoQuant).

Images were acquired with a resolution of 256×256 pixel, zoom 8, a pixel size of 0.1μm and a dwell time of 12.6μs. For each measurement 100 frames were taken and the intensity-weighted mean lifetimes τ (ns) were calculated using PicoQuant SymPhoTime64 software applying a bi-exponential fit. The displayed data were obtained from at least 5 independent experiments.

### Quantitative real time PCR

For qRT-PCR, RNA was isolated from dissected inflorescence meristems using RNeasyMini kit (Qiagen). First strand cDNA was synthesized with 1μg of RNA cDNA using the Superscript III Kit (Invitrogen). Quantitative real-time PCR was performed with 10-fold diluted cDNA using The SsoAdvancedTM Universal SYBR® Green Supermix (Bio-Rad). The mean and standard error were determined using three biological replicates with three technical replicates each. Expression levels were normalized to the references genes TIP41-like (At4g34270) and SAND-domain protein (AT2G28390) (Czechowski *et al*., 2005). Primers used are listed in Tab. 3. Calculation of the relative expression was performed according to Michael W. Pfaffl (Pfaffl, 2001).

### Phenotypic analyses

Rosettes and plants were analyzed by taking photographs using Canon EOS400D camera with an EF-S 60 mm Canon ZOOM lens) of plants growing on soil. Inflorescences were analyzed by taking pictures using stereo microscope (Nikon SMZ25). The inflorescence plastochron was obtained by calculating the average time separating the emergence of 2 successive flower above stage 15 emerging after plant bolting. For SAM area measurement, 30DAG plants growing under LD conditions were used. The primary inflorescence was dissected and imaged using LSM780. The SAM area measurement was done by Fiji.

### Software

Microsoft Word, Excel, and PowerPoint software was used to organize experimental data. Images were analyzed and processed with ImageJ v 1.53c (Schneider *et al*., 2012) and Carl Zeiss ZEN2011. All images were adjusted in “Brightness and Contrast”. VectorNTI (InvitrogenTM) was used for vector maps and sequence analysis. Databank gene research were performed on The Arabidopsis Information Resource (TAIR), (http://www.arabidopsis.org/). Indigo was used for the luciferase assay imaging. Statistical analyses and box plots were realized with GraphPad Prism v 8. For visualization. Using the open-source software MorphoGraphX (MGX) software (https://morphographx.org/) (Barbier de Reuille *et al*., 2015) the surface of the meristem was extracted and the PI signal of the cell wall from layer1 (L1) was projected and used for segmentation of the images to quantify number and size of cells. Cells were segmented manually. MorphographX analysis was performed according to standards defined in the user manual (Barbier de Reuille et al., 2015). The visualisation and counting of nuclei expressing PlaCCI (Desvoyes et al., 2020) was done with Imaris (version 9.1.2, Bitplane, Oxford Instruments plc). Ratios for R2D2 were calculated as described previously (Bhatia *et al*., 2016). All Statistical analyses and data plotting were realized with GraphPad Prism v 8. All images for an experimental set were captured under identical microscope settings.

## Supplemental data

**Suppl Fig. 1:**
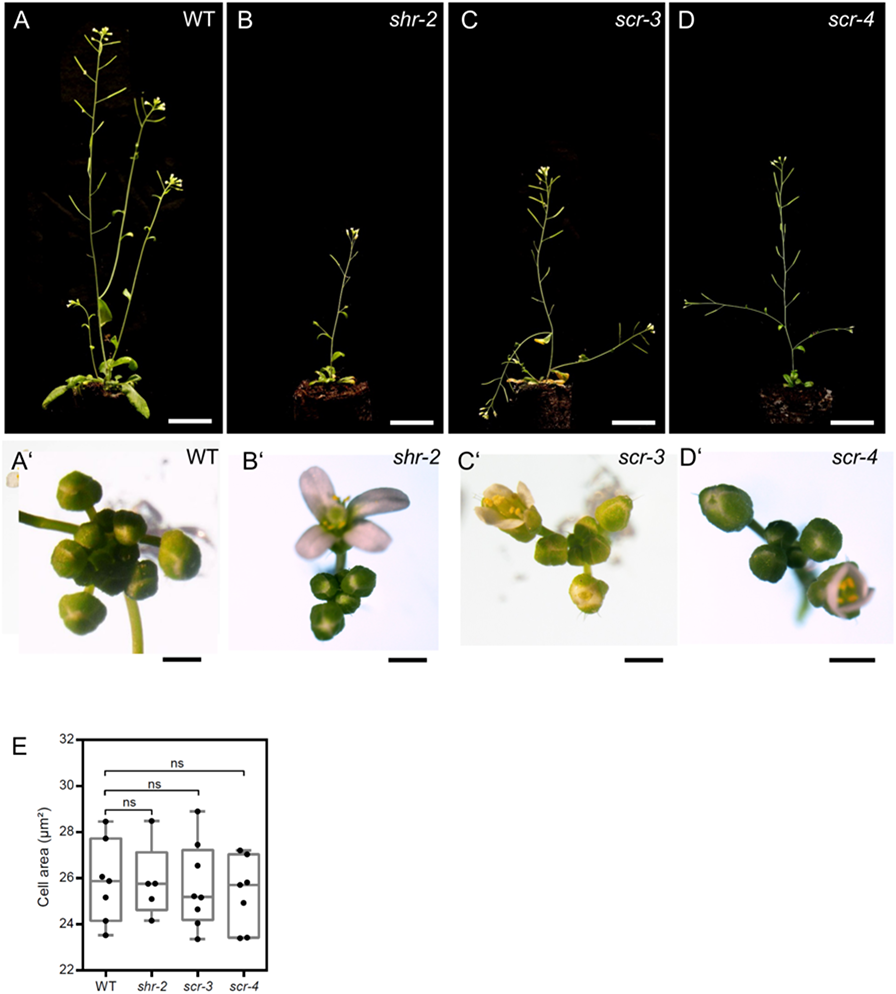
The *shr* and *scr* mutants phenotypes. **(A-D)** Plant phenotypes of 42-day-old WT (n> 30) **(A)**, *shr-2* mutant (n> 35) **(B)**, *scr-3* mutant (n> 30) **(C)** and *scr-4* mutant (n> 30) **(D)**. Scale bar represents 1cm. **(A’-D’)** Top view of 31-day-old inflorescences of WT (n> 30) **(A’)**, *shr-2* mutant (n> 35) **(B’)**, *scr-3* mutant (n> 30) **(C’)** and *scr-4* mutant (n> 30) **(D’)**. Scale bars represent 2 mm. **(E)** Quantification of the epidermal cell area in the meristem region of WT (n=11), *shr-2* mutant (n=4), *scr-3* mutant (n=7) and *scr-4* mutant (n=6). Statistically significant differences were determined by Student’s *t*-test, ns= no significant difference.

**Suppl Fig. 2:**
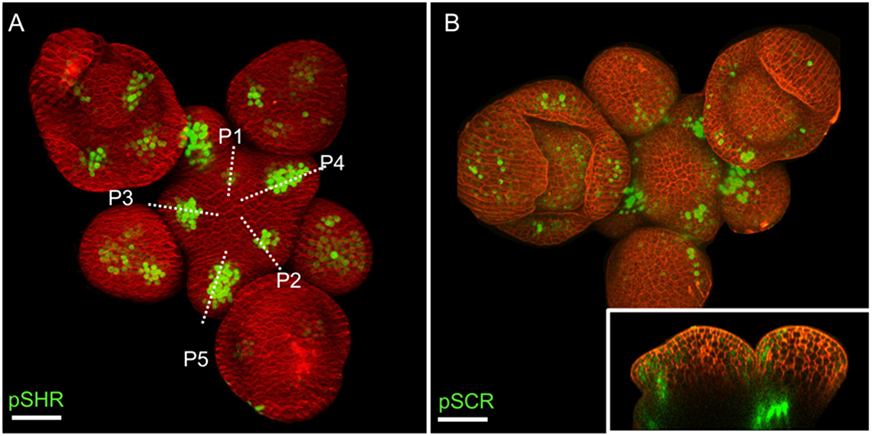
The expression pattern of SHR and SCR in the shoot apical meristem. **(A)** Representative 3D projection of shoot apical meristem at 5 weeks after germination expressing *pSHR:ntdTomato* reporter (green). Cell walls were stained with DAPI (red) (n≥5). Scale bar represents 50 µm. **(B)** Representative 3D projection of shoot apical meristem at 5 weeks after germination expressing *pSCR:H2B-YFP* reporter (green). Cell walls were stained with PI (red) (n≥3). The lower right inset shows an optical longitudinal section through the middle of the SAM. Scale bar represents 50 µm.

**Suppl Fig. 3:**
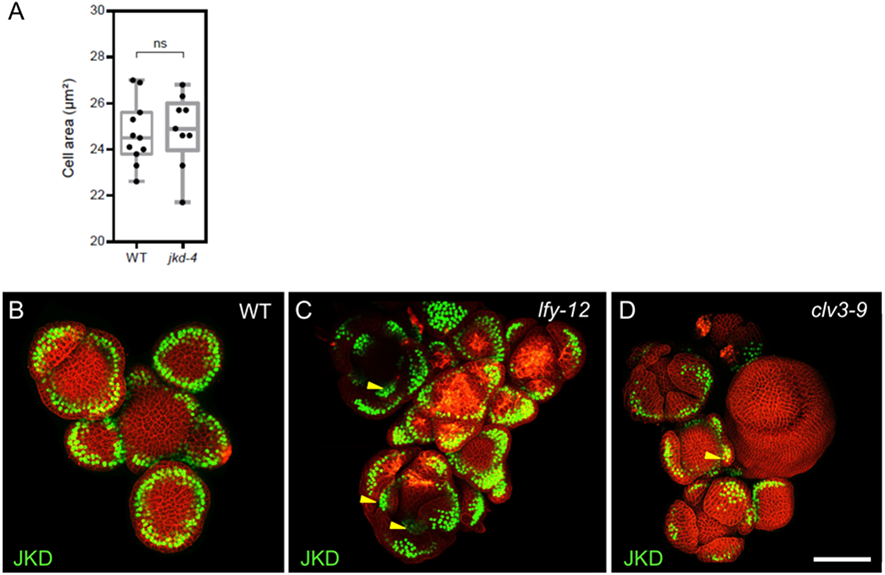
The expression pattern of JKD in *lfy* and *clv3* mutants shoot apical meristem. **(A)** Quantification of the epidermal cell area in the meristem region of WT (n=10) and *jkd-4* mutant (n=10). **(B-D)** Representative 3D projection of shoot apical meristems at 5 weeks after germination expressing *pJKD:JKD-YFP* reporter (green) in WT **(B)**, *lfy-12* mutant **(C)** and *clv3-9* mutant **(D)**. Cell walls were stained with PI (red). Scale bar represents 50 µm. difference. Statistically significant differences were determined by Student’s *t*-test, ns= no significant.

**Suppl Fig. 4:**
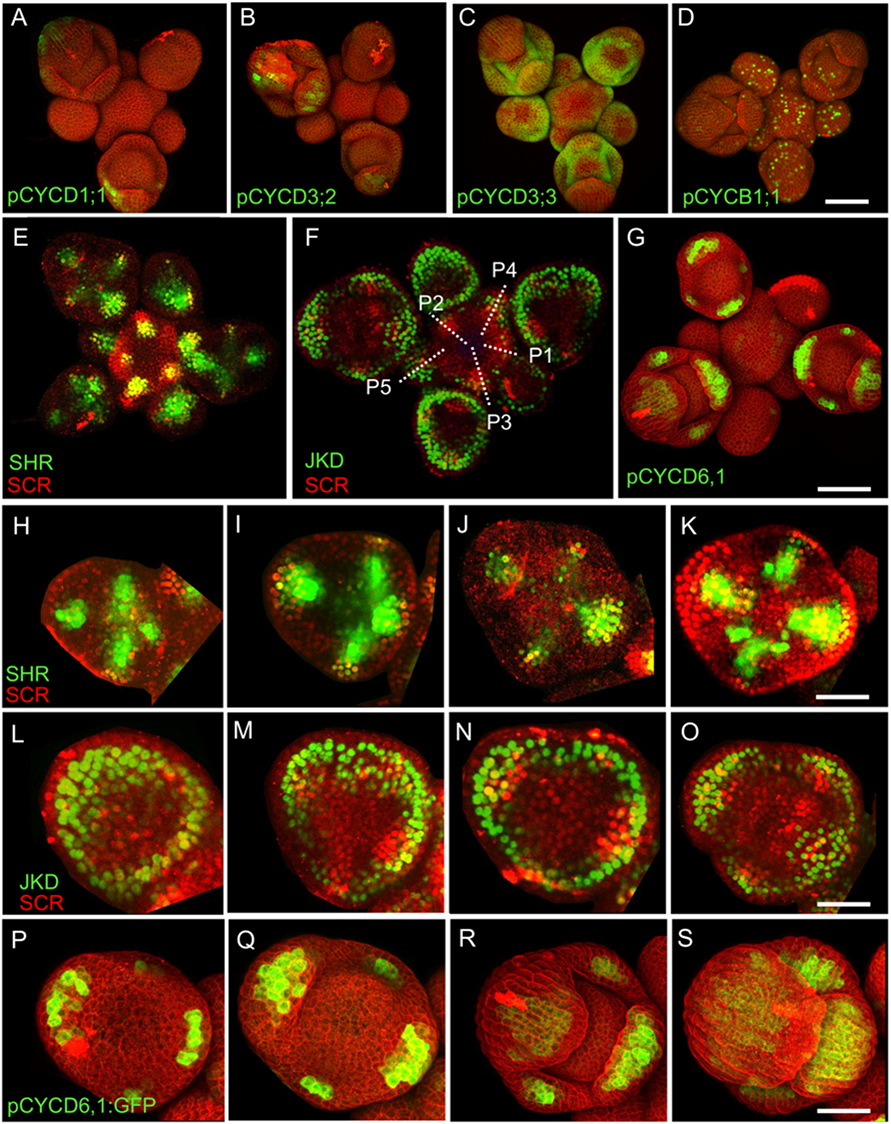
The colocalization of the expression patterns of SHR, SCR, JKD and CYCD6;1 in the shoot apical meristem. **(A-D)** Representative 3D projection of shoot apical meristem at 5 weeks after germination expressing *pCYCD1;1-GFP* reporter (green) **(A)**, *pCYCD3;2-GFP* reporter (green) **(B)**, *pCYCD3;3-GFP* reporter (green) **(C)** and *pCYCB1;1:CYCB1;1-GFP* reporter (green) **(D)**. Cell walls were stained with PI (red). Scale bars, 50 µm. **(E and F)** Representative 3D projection of shoot apical meristems at 5 weeks after germination coexpressing *pSHR:SHR-YFP* reporter (green) and *pSCR:SCR-RFP* reporter (red) (E) and *pJKD:JKD-YFP* reporter (green) and *pSCR:SCR-RFP* reporter (red) (F). Scale bar represents 50 µm. (G) Representative 3D projection of shoot apical meristems expressing *pCYCD6;1:GFP* (green) reporter (n≥5). Cell walls were stained with PI (red). Scale bar represents 50 µm. **(H-S)** Expression of SHR, SCR, JKD and *CYCD6;1* during early stages of flower development. expression is observed at the sepal primordia in the florescence meristem stage 3 **(H, L and P)**, stage 4 **(I, M and Q)**, stage 5 **(J, N and R)** and stage 6 **(K, O and S)**. Scale bar represents 20 µm.

**Suppl Fig. 5:**
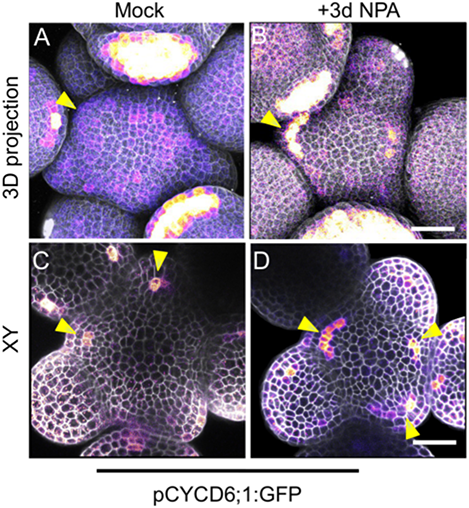
*CYCD6;1* expression responds to auxin. **(A and B)** Representative 3D projection of shoot apical meristems at 5 weeks after germination expressing *pCYCD6;1:GFP* reporter (magenta) three days after mock (n≥3) **(A)** or 100 µM NPA treatment (n≥6) **(B)**. **(C and D)** Transversal optical sections of **(A)** and **(B)** respectively. Yellow arrowheads in **(A)**, **(B)**, **(C)** and **(D)** show primordia with pCYCD6;1:GFP expression. Cell walls were stained with DAPI (gray). Scale bar represents 50 µm.

**Suppl Fig. 6:**
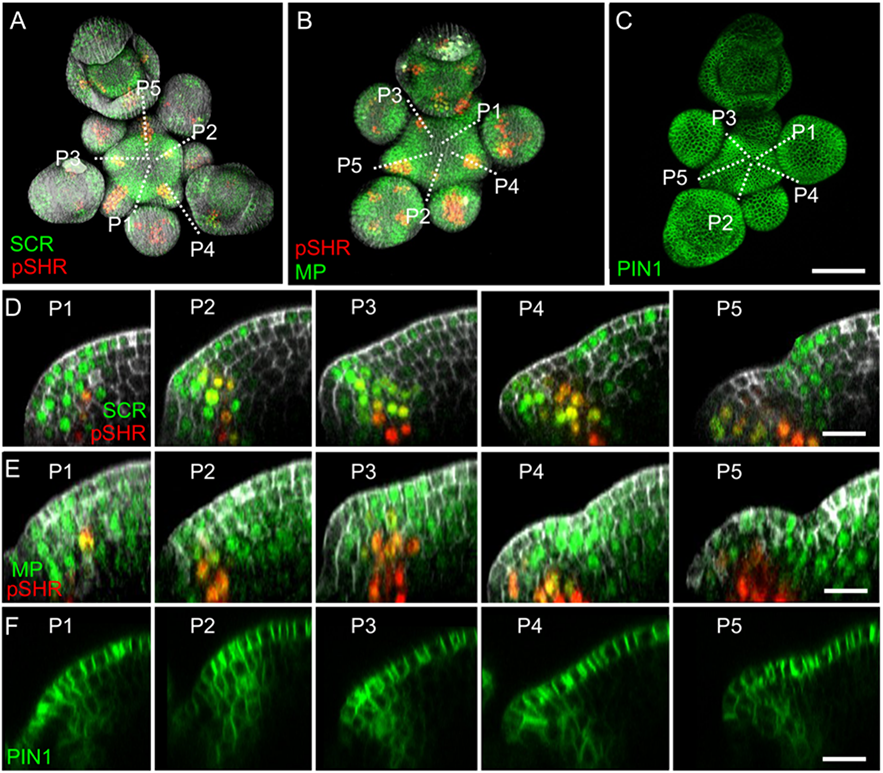
The expression patterns of SHR, SCR, MP and PIN1 in the shoot apical meristem. **(A and B)** Representative 3D projection of shoot apical meristems at 5 weeks after germination coexpressing *pSCR:SCR-YFP* reporter (green) and *pSHR:ntdTomato* reporter (red) **(A)** and the *pMP:MP-GFP* (green) and *pSHR:ntdTomato* (red) reporters **(B)**. Cell walls were stained with DAPI (gray). Scale bar represents 50 µm. **(C)** Representative 3D projection of shoot apical meristems at 5 weeks after germination expressing *pPIN1:PIN1-GFP* reporter (green). Scale bar represents 50 µm. **(D-F)** Longitudinal optical sections through the middle of five successive primordia (representative section orientation shown by dotted line in **(A)**, **(B)** and **(C)** respectively). Scale bar represents 20 µm.

**Suppl Fig. 7:**
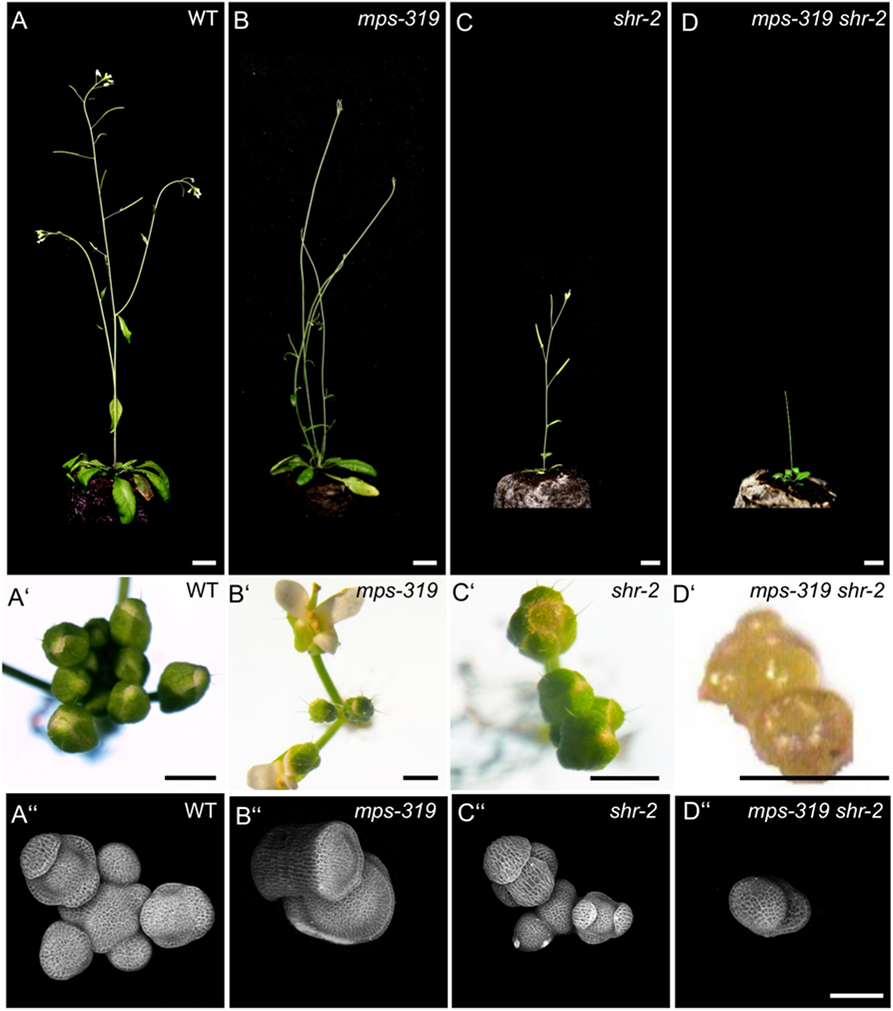
*shr* mutant and *shr mps-319* double-mutant phenotypes. **(A-D)** Plant phenotypes of 42-day-old WT **(A)**, *mps-319* mutant **(B)**, *shr-2* mutant **(C)** and *mps-319 shr-2* double mutant **(D)**. Scale bar represents 1 cm. **(A’-D’)** Top view of 31-day-old inflorescences of WT **(A’)**, *mps-319* mutant **(B’)**, *shr-2* mutant **(C’)** and *mps-319 shr-2* double mutant **(D’)**. Scale bars represent 2 mm. **(A’’-D’’)**. Representative 3D projection of shoot apical meristems at 5 weeks after germination from WT **(A’’)**, *mps-319* mutant **(B’’)**, *shr-2* mutant **(C’’)** and *mps-319 shr-2* double mutant **(D’’)**. Scale bar represents 50 µm.

**Suppl Fig. 8:**
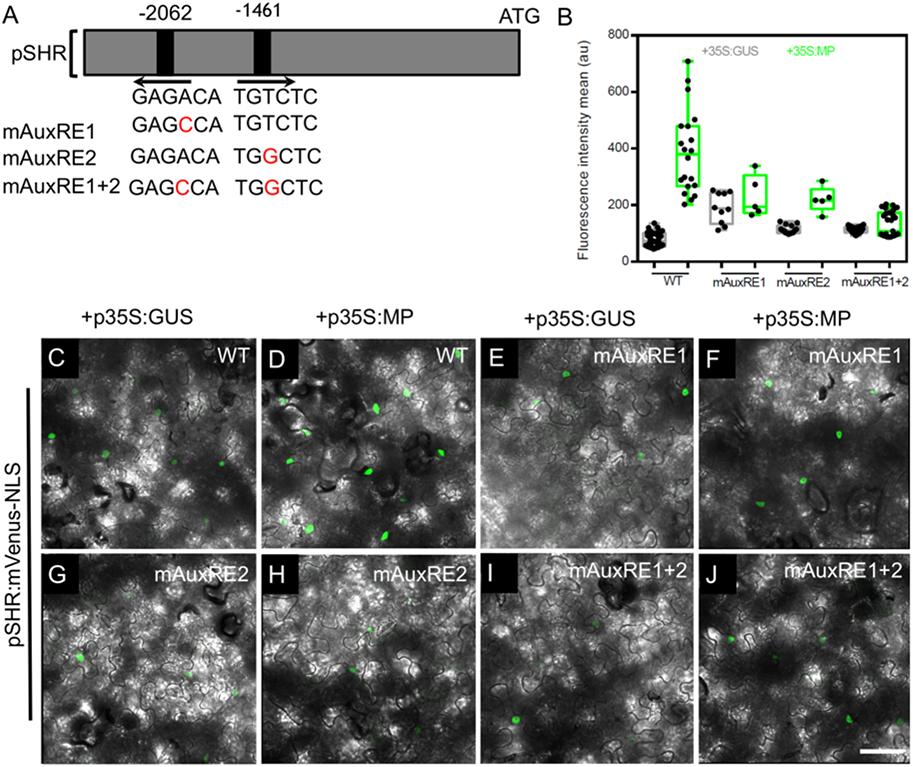
MP Regulates *SHR* expression in the shoot apical meristem. **(A)** Schematic representation of the *SHR* promoter. The positions of two auxin response elements are shown. Overview of mutated promoter versions of *pSHR*. AuxREs were mutated and multiple combinations of these mutated motifs were combined into a single promoter. The original AuxRE sequence GAGACA was mutated to GAGCCA (mAuxRE1), the original AuxRE sequence TGTCTC was mutated to TGGCTC (mAuxRE2). **(B)** Quantification of mVenus fluorescence signal intensity from leaves transiently transformed with mVenus-NLS under the control of the wild-type SHR promoter or under the control of the SHR promoter with mutations in AuxRE motifs in the presence of *p35S:GUS* or *p35S:MP*. **(C-J)** Leaves transiently transformed with mVenus-NLS under the control of the wild-type SHR promoter **(C and D)** or under the control of the SHR promoter with mutations in AuxRE motifs, mAuxRE1 **(E and F)** mAuxRE2 **(G and H)** and mAuxRE1+2 **(I and J)** together with *p35S:MP* or *p35S:GUS*. mVenus fluorescence signals were detected by confocal microscopy. Scale bar represents 50 µm.

**Suppl Fig. 9:**
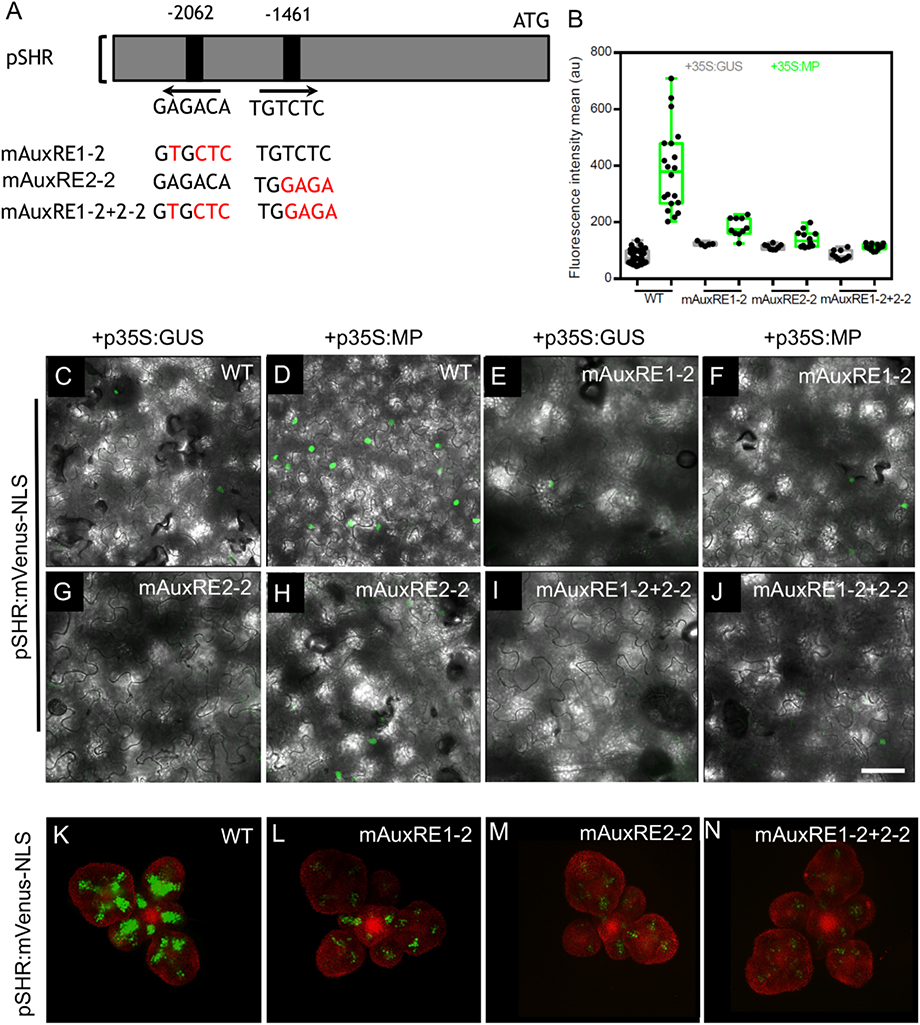
MP Regulates *SHR* expression in the shoot apical meristem. **(A)** Schematic representation of the *SHR* promoter. The positions of two auxin response elements are shown. Overview of mutated promoter versions of *pSHR*. AuxREs were mutated and multiple combinations of these mutated motifs were combined into a single promoter. The original AuxRE sequence GAGACA was mutated to GTGCTC (mAuxRE1), the original AuxRE sequence TGTCTC was mutated to TGGAGA (mAuxRE2-2). **(B)** Quantification of mVenus fluorescence signal intensity from leaves transiently transformed with mVenus-NLS under the control of the wild-type SHR promoter or under the control of the SHR promoter with mutations in AuxRE motifs in the presence of *p35S:GUS* or *p35S:MP*. **(C-J)** Leaves transiently transformed with mVenus-NLS under the control of the wild-type SHR promoter **(C and D)** or under the control of the SHR promoter with mutations in AuxRE motifs, mAuxRE1-2 **(E and F)** mAuxRE2-2 **(G and H)** and mAuxRE1-2+2-2 **(I and J)** together with *p35S:MP* or *p35S:GUS*. mVenus fluorescence signals were detected by confocal microscopy. Scale bar represents 50 µm. **(K–N)** Representative 3D projection of shoot apical meristems at 5 weeks after germination expressing mVenus-NLS under the control of the wild-type SHR promoter **(K)**, and under the control of the SHR promoter with mutations in AuxRE motifs mAuxRE1-2 **(L)** mAuxRE2-2 **(M)** and mAuxRE1-2+2-2 **(N)**; Chlorophyll (red). Scale bar represents 50 µm.

**Suppl Fig. 10:**
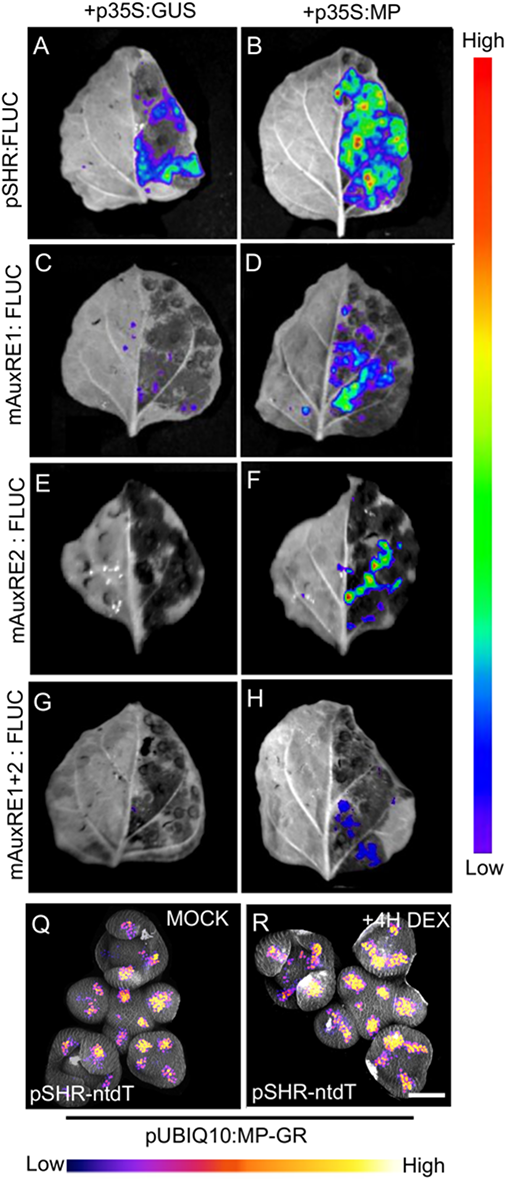
MP induces the expression of SHR *in planta*. **(A-H)** Representative images of leaves transformed with pSHR variants (pSHR (A and B), mAuxRE1 **(C and D)**, mAuxRE2 **(E and F)** and mAuxRE1+2 **(G and H)** draving the reporter gene firefly luciferase (FLUC) together with *p35S:MP* or *p35S:GUS*. D-luciferin was used as the substrate of FLUC. No luminescence could be detected without the substrate D-Luciferin. Color code indicate relative signal intensities (red: high; violet: low). **(Q and R)** Representative 3D projection of shoot apical meristems at 5 weeks after germination expressing *pSHR:ntdTomato* reporter (magenta) in *pUBIQ10:MP-GR* plants 4 hr after mock **(Q)** (n = 4) and 4 hr after DEX treatment **(P)** (n = 4). Cell walls were stained with DAPI (gray). Scale bar represents 50 µm.

**Suppl Fig. 11:**
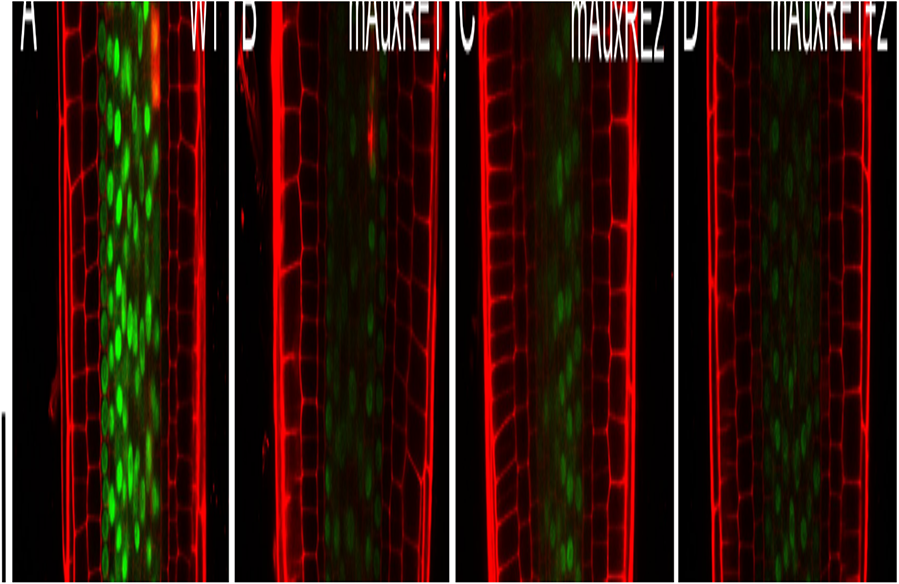
The expression pattern of the different promoter versions of *SHR* in the root apical meristem. **(A–G)** Representative images of root apical meristem at 5 days after germination expressing mVenus-NLS driven by the wild-type SHR promoter **(A)**, and the SHR promoter with mutations in AuxRE motifs mAuxRE1 **(B)**, mAuxRE2 **(C)**, mAuxRE1+2 **(D)**, mAuxRE1-2 **(E)**, mAuxRE2-2 **(F)** and mAuxRE1-2+2-2 **(G)**. **(H)** Representative images of 5-d-old root tips that coexpressing the *pMP:MP-GFP* reporter (green) and *pSHR:ntdTomato* reporter (red). Cell walls were stained with PI (red). Scale bar represents 50 µm.

**Suppl Fig. 12:**
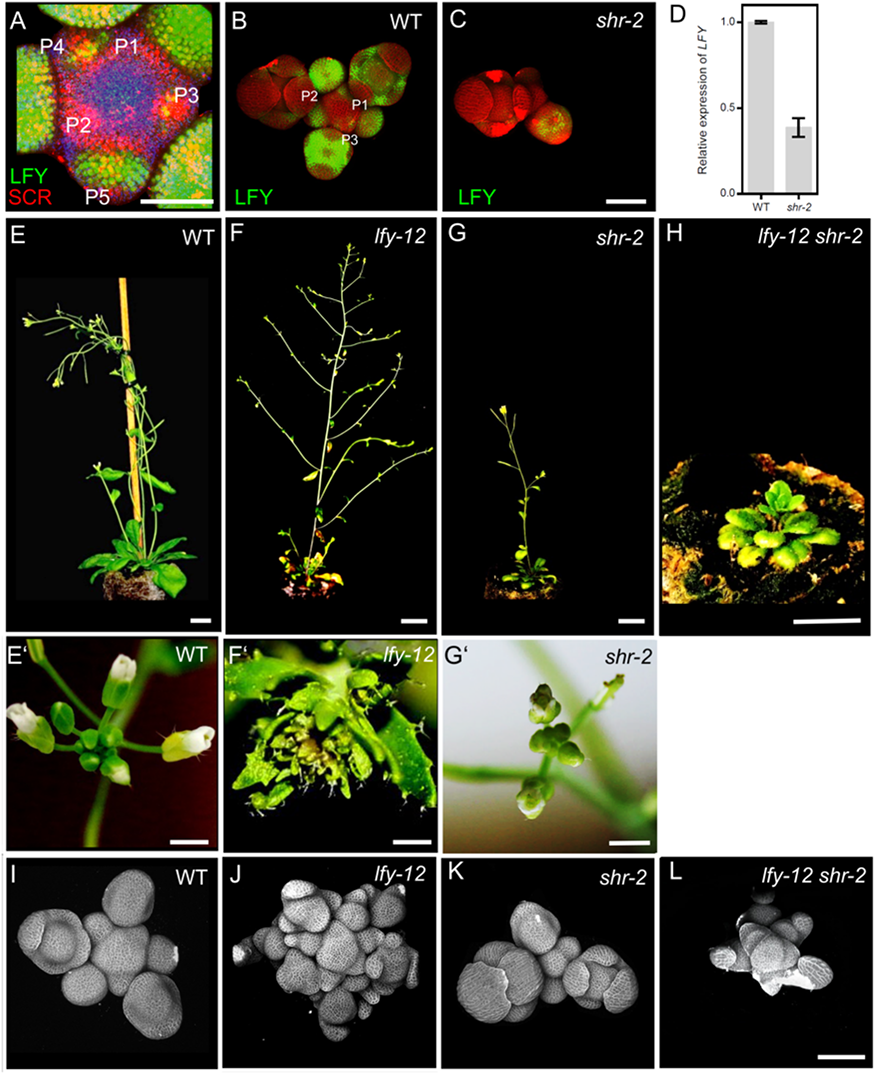
*LFY* act downstream of SHR in the shoot apical meristem. **(A)** Representative 3D projection of shoot apical meristems at 5 weeks after germination coexpressing *pLFY:LFY-GFP* (green) and *pSCR:SCR-RFP* (red) reporters. Chlorophyll (blue). Scale bar represents 50 µm. **(B and C)** Representative 3D projection of shoot apical meristems expressing *pLFY:LFY-GFP* reporter (green) in WT(n≥3) **(L)** and *shr-2* mutant (n≥3) **(M)**. Cell walls were stained with PI (red). Scale bar represents 50 µm. **(D)** Quantitative real-time PCR analysis showing the relative expression levels of *LFY* in WT and *shr-2* mutant shoot apical meristems. The expression level in Col-0 is set to 1 and error bars show standard deviation. Expression levels were normalized using At4g34270 and AT2G28390. **(E-H)** Plant phenotypes of 45-day-old WT **(E)**, *lfy-12* mutant **(F)**, *shr-2* mutant **(G)** and *lfy-12 shr-2* double mutant **(H)**. Scale bar represents 1 cm. **(E’-G’)** Top view of 31-day-old inflorescences of WT **(E’)**, *lfy-12* **(F’)** mutant and *shr-2* mutant **(G’)**. Scale bar represents 2 mm. **(I-L)** Representative 3D constraction of shoot apical meristems at 5 weeks after germination from WT **(I)**, *lfy-12* mutant **(J)**, *shr-2* mutant **(K)** and *lfy-12 shr-2* double mutant **(L)**. Scale bar represents 50 µm.

**Suppl Fig. 13:**
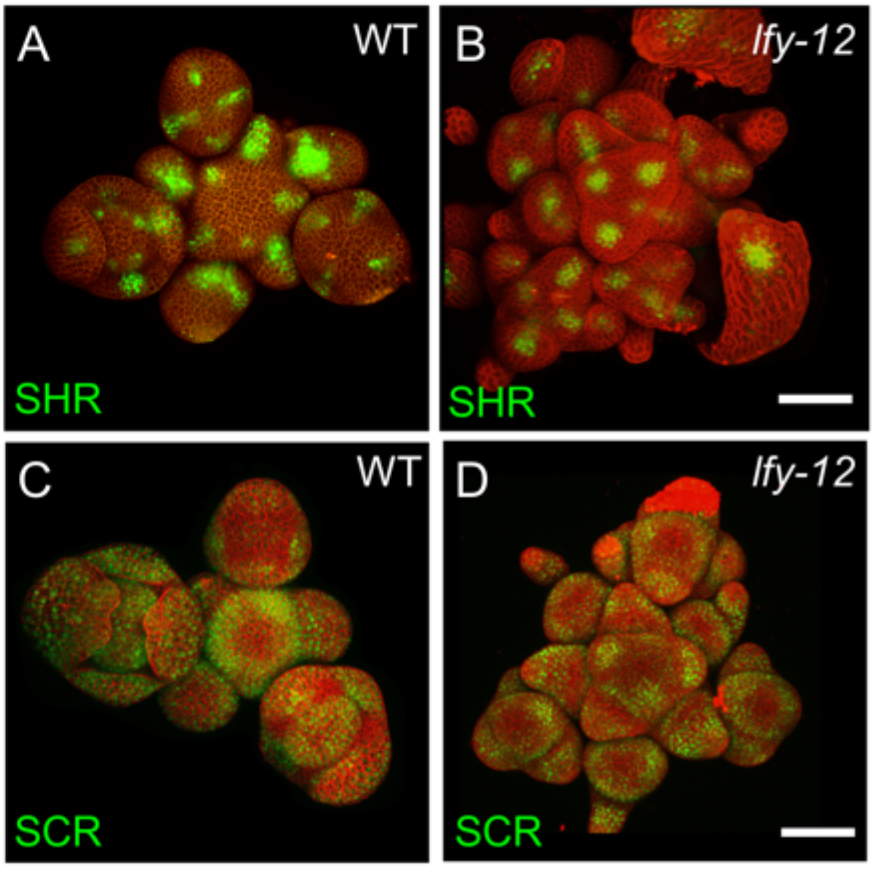
The expression pattern of SHR and SCR in *lfy* mutant shoot apical meristem. **(A and B)** Representative 3D projection of shoot apical meristems at 5 weeks after germination expressing *pSHR:SHR-YFP* reporter (green) in WT **(A)** and *lfy-12* mutant **(B)**. Cell walls were stained with PI (red). Scale bar represents 50 µm. **(C and D)** Representative 3D projection of shoot apical meristems expressing *pSCR:SCR-YFP* reporter (green) in WT **(C)** and *lfy-12* mutant **(D)**. Cell walls were stained with PI (red). Scale bar represents 50 µm.

**Suppl Fig. 14:**
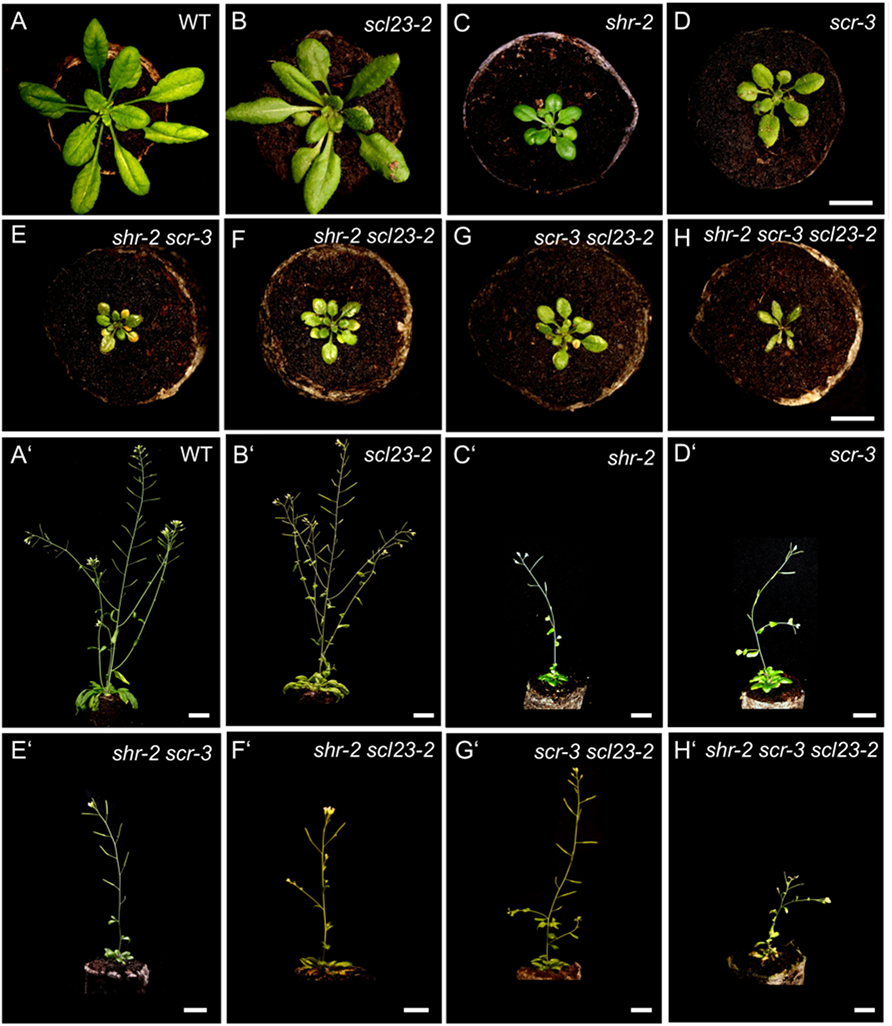
Genetic combinations of *shr*, *scr* and *scl23*. **(A-H)** Top view of 21-day-old rosettes of WT **(A)**, *scl23-2* mutant **(B)**, *shr-2* mutant **(C)**, *scr-3* mutant **(D)**, *scr-3 shr-2* double mutant **(E)**, *scl23-2 shr-2* double mutant **(F)**, *scl23-2 scr-3* double mutant **(G)**, *scr-3 shr-2 scl23-2* triple mutant **(H)**. Scale bar represents 1 cm. **(A’-H’)** Plant phenotypes of 6 weeks old WT **(A’)**, *scl23-2* mutant **(B’)**, *shr-2* mutant **(C’)**, *scr-3* mutant **(D’)**, *scr-3 shr-2* double mutant **(E’)**, *scl23-2 shr-2* double mutant **(F’)**, *scl23-2 scr-3* double mutant **(G’)**, *scr-3 shr-2 scl23-2* triple mutant **(H’)**. Scale bar represents: 2 cm **(A’ and B’)**, 1 cm **(C’-H’)**.

**Suppl Fig. 15:**
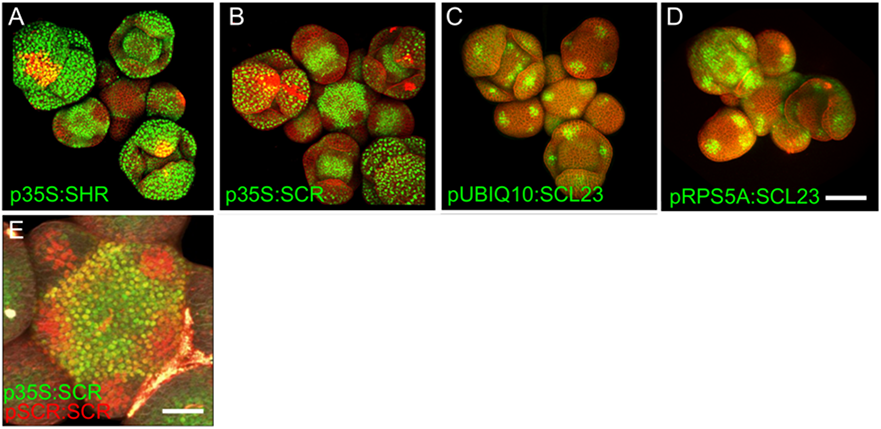
The overexpression of SHR, SCR and SCL23 in the SAM. **(A-D)** Representative 3D projection of shoot apical meristems at 5 weeks after germination expressing *p35S:SHR-GFP* reporter (green) **(A)**, *p35S:SCR-GFP* reporter (green) **(B)**, *pUBIQ10:SCL23-mVenus* reporter (green) **(C)** and *pRPS5A:SCL23-mVenus* reporter (green) **(D)**. Cell walls were stained with PI (red). Scale bar represents 50 µm. **(E)** Top view of the inflorescence apex coexpressing the *p35S:SCR-GFP* reporter (green) and the *pSCR:SCR-RFP* reporter (red). Cell walls were stained with DAPI (gray). Scale bar represents 50 µm.

**Suppl Fig. 16:**
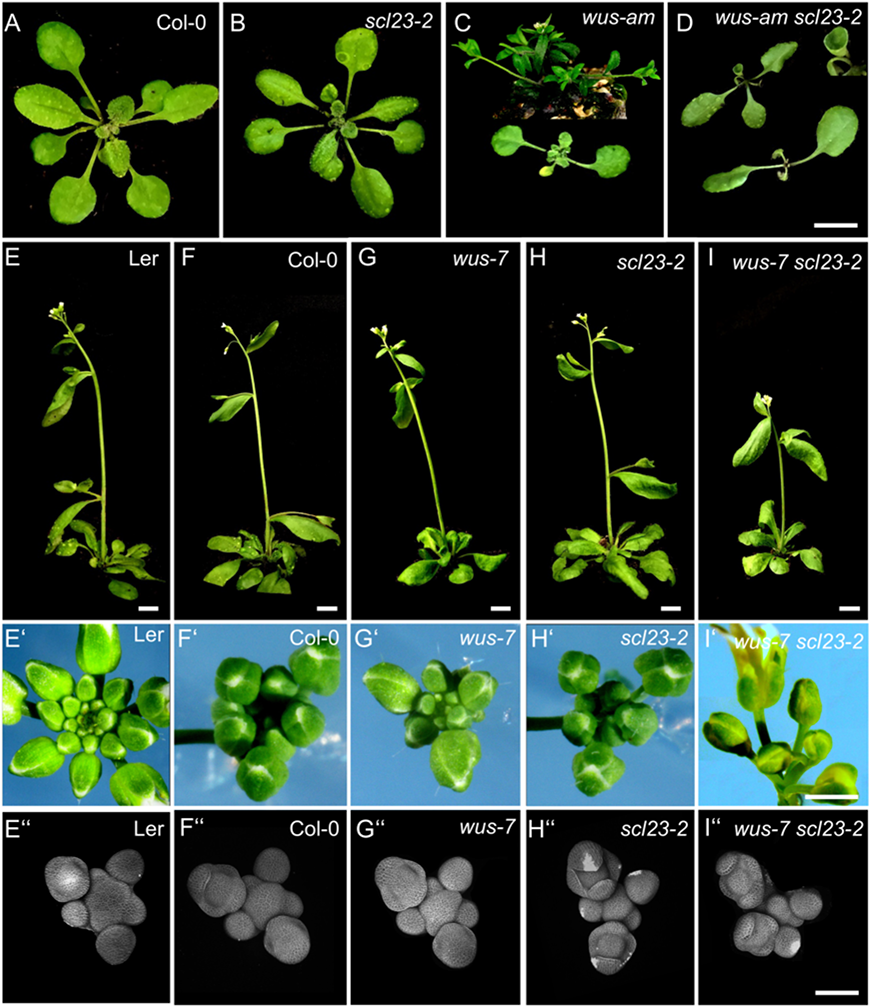
WUS and SCL23 cooperatively control shoot stem cell homeostasis in the shoot apical meristem. **(A-D)** Top view of 21-day-old rosettes of Col-0 **(A)**, *scl23-2* mutant **(B)**, *wus-am* mutant **(C)** and *scl23-2 wus-am* double mutant **(D)**. Scale bar represents 1 cm. **(E-I)** plant phenotypes of 36 old L.er **(E)**, Col-0 **(F)**, *scl23-2* mutant **(G)**, *wus-7* mutant **(H)** and *scl23-2 wus-7* double mutant (I). Scale bar represents 1 cm. **(E’-I’)** Top view of 35-day-old inflorescences of Ler **(E’)**, Col-0(F’), *scl23-2* mutant **(G’)**, *wus-7* mutant **(H’)** and *scl23-2 wus-7* double mutant **(I’)**. Scale bar represents 2 mm. **(E’’-I’’)** Representative 3D projection of shoot apical meristems at 5 weeks after germination from Ler **(E’’)**, Col-0 **(F’’)**, *scl23-2* mutant **(G’’)**, *wus-7* mutant **(H’’)** and *scl23-2 wus-7* double mutant (I’’). Cell walls were stained with PI (gray). Scale bar represents 50 µm. **(A-D)** Top view of 21-day-old rosettes of Col-0 **(A)**, *scl23-2* mutant **(B)**, *wus-am* mutant **(C)** and *scl23-2 wus-am* double mutant **(D)**. Scale bar represents 1 cm. **(E-I)** plant phenotypes of 36 old L.er **(E)**, Col-0 **(F)**, *scl23-2* mutant **(G)**, *wus-7* mutant **(H)** and *scl23-2 wus-7* double mutant **(I)**. Scale bar represents 1 cm. **(E’-I’)** Top view of 35-day-old inflorescences of Ler **(E’)**, Col-0(F’), *scl23-2* mutant **(G’)**, *wus-7* mutant **(H’)** and *scl23-2 wus-7* double mutant **(I’)**. Scale bar represents 2 mm. **(E’’-I’’)** Representative 3D projection of shoot apical meristems at 5 weeks after germination from Ler **(E’’)**, Col-0 **(F’’)**, *scl23-2* mutant **(G’’)**, *wus-7* mutant **(H’’)** and *scl23-2 wus-7* double mutant (I’’). Cell walls were stained with PI (gray). Scale bar represents 50 µm

**Suppl Fig. 17:**
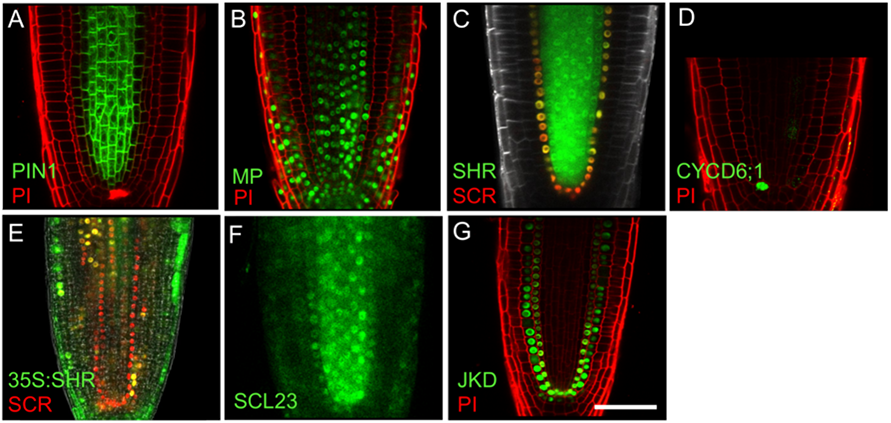
The overlapping expression pattern of PIN1, MP, SHR, SCR, *CYCD6;1*, SCL23 and JKD in the RAM. **(A-D)** Representative images of root apical meristem at 5 days after germination expressing, *pPIN1:PIN1-GFP* reporter (green) **(A)**, *pMP:MP-GFP* reporter (green) **(B)**, *pSHR:YFP-SHR* reporter (green) and *pSCR:SCR-RFP* reporters (red) **(C)**, *pCYCD6;1:GFP* reporter (green) **(D)**, *p35S:SHR-GFP* reporter (green) and *pSCR:SCR-RFP* reporter (red) **(E)**, *pSCL23:SCL23-YFP* reporter (green) **(F)** and *pJKD:JKD-YFP* reporter (green) **(G)**. Cell walls were stained with PI (red) in **(A) (B) (D) (G)** or DAPI (gray) in **(C)**. Scale bar represents 50 µm.

**Suppl Fig. 18:**
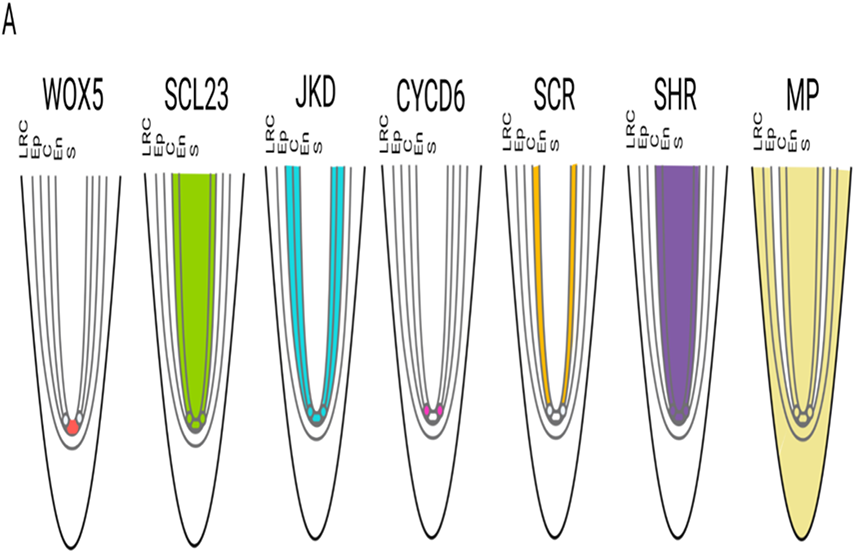
Proposed model for SHR-SCR-SCL23-JKD regulatory network function in the RAM. **(A)** Schematic representation of observed expression patterns of *WOX5*, *SCL23*, *JKD, CYCD6, SCR, SHR* and *MP* showing typical expression domains in the RAM. **(B)** Schematic molecular models for SHR-SCR-SCL23-JKD gene regulatory network function in the RAM, genes with genetic and/or biochemical interactions are indicated. Lines with arrows depict positive regulation, and with bars depict negative regulation. Dashed lines with arrows depict intercellular protein movement. Overlap between circles describe protein–protein interactions. In the RAM, *SHR* is transcribed in the stele (S) in response to auxin-dependent activation of MP, and SHR protein moves to the CEI, where SHR activates *SCR*, *SCL23* and *JKD* expression. The multimeric protein complexes then induce formative division by activating expression of *CYCD6*. High levels of auxin in the CEI also contribute to *CYCD6* activation. In the QC, the SHR-SCR-SCL23-JKD protein complex positively regulates expression of *WOX5*. CEI= Cortex/endodermis initial, S=Stele, QC=Quiescent center.

## List of Figures

Fig. 10: The *shr* and *scr* mutant phenotypes

Fig. 11: SHR and SCR functions modulate auxin signalling in the shoot apical meristem

Fig. 12: Expression patterns of SHR and SCR in the shoot apical meristem

Fig. 13: *In vivo* FRET–FLIM quantification of SHR-SCR association in the SAM

Fig. 14: SHR regulates *SCR* expression in the SAM

Fig. 15: JKD functions and expression pattern in the SAM

Fig. 16: SHR regulates *CYCD6;1* expression in the SAM

Fig. 17: SHR and SCR expressions respond to auxin

Fig. 18: SHR and SCR act downstream of MP in the SAM

Fig. 10: MP Regulates *SHR* expression in the SAM

Fig. 11: SCL23, SCR and SHR proteins show spatially different patterns but perform similar functions in the SAM

Fig. 12: SCL23 and WUS physically interact and are negatively regulated by CLV signalling

Fig. 13: Proposed model for SHR-SCR-SCL23-JKD regulatory network function in the SAM

## List of Supplementary Figures

Suppl Fig. 7: The *shr* and *scr* mutants phenotypes

Suppl Fig. 8: The expression pattern of SHR and SCR in the shoot apical meristem

Suppl Fig. 9: The expression pattern of JKD in *lfy* and *clv3* mutants shoot apical meristem

Suppl Fig. 10: The colocalization of the expression patterns of SHR, SCR, JKD and CYCD6;1 in the shoot apical meristem

Suppl Fig. 11: *CYCD6;1* expression responds to auxin

Suppl Fig. 12: The expression patterns of SHR, SCR, MP and PIN1 in the shoot apical meristem

Suppl Fig. 7: *shr* mutant and *shr mps-319* double-mutant phenotypes

Suppl Fig. 8: MP Regulates *SHR* expression in the shoot apical meristem

Suppl Fig. 9: MP Regulates *SHR* expression in the shoot apical meristem

Suppl Fig. 10: MP induces the expression of SHR *in planta*

Suppl Fig. 11: The expression pattern of the different promoter versions of *SHR* in the root apical meristem

Suppl Fig. 12: *LFY* act downstream of SHR in the shoot apical meristem

Suppl Fig. 13: The expression pattern of SHR and SCR in *lfy* mutant shoot apical meristem

Suppl Fig. 14: Genetic combinations of *shr*, *scr* and *scl23*

Suppl Fig. 15: The overexpression of SHR, SCR and SCL23 in the SAM

Suppl Fig. 16: WUS and SCL23 cooperatively control shoot stem cell homeostasis in the shoot apical meristem

Suppl Fig. 17: The overlapping expression pattern of PIN1, MP, SHR, SCR, *CYCD6;1*, SCL23 and JKD in the RAM

Suppl Fig. 18: Proposed model for SHR-SCR-SCL23-JKD regulatory network function in the RAM

## List of Tables

Tab. 10: Enzymes used in this study

Tab. 11: Primers used for cloning

Tab. 12: Primers used for qRT-PCR

Tab. 13: Primers used for genotyping

Tab. 14: Entry plasmids used in this study

Tab. 15: Destination plasmids used in this study

Tab. 16: Destination plasmids used for luciferase assay

Tab. 17: Mutants used in this study

Tab. 18: Transgenic lines used in this study

